# Deep dynamical models of single-cell multiomic velocities predict loss-of-function and rescue perturbations in B cells

**DOI:** 10.1101/2025.04.24.650458

**Authors:** Alireza Karbalayghareh, Benedikt Pelzer, Christopher R. Chin, Ari Melnick, Darko Barisic, Christina S. Leslie

**Affiliations:** Computational and Systems Biology Program, Memorial Sloan Kettering Cancer Center, New York, NY 10065, USA; Department of Medicine, Weill Cornell Medicine, Cornell University, New York, NY 10065, USA; Meyer Cancer Center, Weill Cornell Medical College, New York, NY 10065, USA

## Abstract

We present DynaVelo, a generative neural ordinary differential equation model that learns the joint dynamics of gene expression and transcription factor (TF) motif activities in evolving cell systems using single-cell multiome with joint gene expression and chromatin accesibility readout. DynaVelo leverages partial RNA velocity information together with single-cell TF motif accessibility data to improve the modeling of cell state dynamics and identification of TF drivers. We show that DynaVelo recovers the complex and bifurcating *in vivo* dynamics of wildtype murine germinal center (GC) B cells and reveals how these cell dynamics change under loss-of-function mutations in epigenetic regulators *Arid1a* and *Ctcf*. DynaVelo resolves how TF motif activities evolve along latent time trajectories using analysis of training cells or through generated trajectories from the model. *In silico* perturbation analysis further enables DynaVelo to infer dynamic and cell-state-specific gene regulatory networks (GRNs), recovering many known TF-to-gene edges in the wildtype GC GRN and predicting those that are disrupted in mutants. Finally, *in silico* gene and TF perturbations allow both the prediction of cell dynamics under loss-of-function genetic mutations and the identification of TF perturbations to rescue loss-of-function dynamic and immunological phenotypes. This analysis predicted that *Ctcf* knockout would rescue *Arid1a* loss-of-function phenotype in the GC reaction and nominated *Bcl6* and *Stat3* as additional TFs whose knockout would rescue *Arid1a* loss. We validated these predictions *in vivo* using double heterozygous mutant mice, confirming rescue of the *Arid1a* dark zone phenotype in all cases and quantitatively assessing model predictions using multiome in *Arid1aHet;CtcfHet* double heterozygous mice. DynaVelo therefore provides a powerful new deep learning framework for modeling and perturbing dynamic cell systems by harnessing single-cell multiome data sets.

## Introduction

Recent single-cell technologies have succeeded in measuring multiple genome-wide readouts from the same individual cells, often pairing the transcriptome with epigenomic measurements such as chromatin accessibility, histone modifications, TF occupancy, or the 3D chromatin interactome^1–14^. These single-cell readouts profile the phenotypic and epigenetic states of cells at a single snapshot of time. However, cells are intrinsically dynamic systems where complex regulatory mechanisms govern the cellular gene expression and epigenetic programs that determine cell fate over time. Since real-time *in vivo* measurements of these modalities are generally not feasible, computational methods have been developed to infer the dynamics of cells from single-snapshot datasets. In these approaches, single cells from the same experimental timepoint are viewed as observations sampled from a dynamically evolving system, and each cell is assigned an inferred pseudotime or latent time relative to this presumed dynamic process. One prominent idea to emerge from these efforts was the concept of RNA velocity, initially proposed by Velocyto^15^ and later enhanced by scVelo^16^, which models spliced and unspliced reads from scRNA-seq using ordinary differential equations (ODEs) to estimate the rate of change of gene expression in an implicit dynamic process. In particular, RNA velocities are useful for providing pseudotemporal *directional* information in inferring gene expression trajectories, although velocities are only computable for a subset of genes and may lack robustness^17,18^. To date, single-cell multiome data – providing both transcriptomic and chromatin accessibility readouts from the same cells (scRNA+ATAC-seq) – have not been exploited in velocity-based methods for inferring cell dynamics. Importantly, scATAC-seq allows the computation of single-cell TF binding motif accessibilities, which can serve as a proxy for the regulatory activity of the corresponding TFs. Therefore, these TF motif accessibilities and their underlying velocities should provide information beyond RNA expression and RNA velocities and enable improved modeling of dynamic cell state changes.

Here we present a deep learning model called DynaVelo for inferring cell dynamics from single-cell multiome data that combines both RNA and TF motif accessibility modalities and predicts shifts in cell-fate trajectories under genetic perturbations. Our main hypothesis is that these two modalities influence each other in dynamic cellular systems: the accessibility of TF motifs, as proxies for TF regulatory activity, affects the RNA expression level of target genes, while the RNA expression of TFs, their cofactors, and other epigenetic regulators modulates downstream changes in chromatin state. DynaVelo trains a generative neural ODE to jointly model the dynamics of both gene expression and TF motif accessibility at a single-cell level, moving beyond RNA velocity methods like scVelo and an analogous velocity method for multiome, MultiVelo^19^, as well as recent machine learning approaches like Dynamo^20^ and scDiffEq^21^ that model the dynamics of gene expression alone.

We reasoned that DynaVelo’s use of single-cell TF motif activity data would be especially relevant in cellular systems with rapid and complex dynamics where cell state transitions are driven by TFs. Furthermore, to be able to validate our findings, we used a setting where loss of function of epigenetic regulators is known to alter these TF activities and cell fate decisions. We therefore applied DynaVelo to murine germinal center (GC) B cells representing three genotypes: in the wildtype (WT) setting and in GC B cells bearing heterozygous loss-of-function mutations in Arid1a (*Arid1aHet*), a subunit of the canonical BAF chromatin remodeling complex that is commonly somatically altered in human B cell lymphomas^22,23^, and Ctcf (*CtcfHet*), a key factor involved in 3D chromatin organization^24,25^. Here we found that DynaVelo can recapitulate the complex *in vivo* dynamics of the GC reaction, generate cells and trajectories *de novo* from the model, and recover evolving TF activities and a dynamic gene regulatory network (GRN) over latent time. Importantly, DynaVelo can both predict the impact of loss-of-function genetic perturbations and identify TF perturbations that rescue loss-of-function dynamics and immunological phenotypes. Thus, DynaVelo provides a powerful generative AI framework for modeling and perturbing dynamic cellular systems from single-snapshot single-cell multiome data.

## Results

### DynaVelo learns a latent neural ODE model for single-cell dynamics from multiomic data

DynaVelo is a neural ODE framework for learning cellular dynamics from single-cell multiome data, where paired scATAC-seq and scRNA-seq readouts are available in each cell. As TF motif accessibility is an interpretable cell-level readout derived from chromatin accessibility, we chose first to use chromVAR^26^ to obtain a cell-by-TF-activity matrix from the scATAC readout. We also obtain a cell-by-gene matrix from scRNA readout after preprocessing, normalization, and filtering out unused genes (**Methods**). DynaVelo learns to model the joint dynamics of RNA expression and TF motif accessibility in each cell. Since solving these ODEs in the original high dimensional input spaces could be challenging, we first embed the cells, using both the gene expression and TF motif accessibility inputs, in a lower dimensional latent space where we use neural ODEs to model the dynamics of cells (**Fig. 1a**). The mapping of cells to the latent space is performed by a variational encoder. Single-cell states and velocities learned in the latent space define a vector field over cells, and these values are mapped again to the observational space by a decoder. In addition to the latent states and velocities, DynaVelo also learns latent times for all cells, providing an ordering of cells in the joint vector field. Similar to variational autoencoders, DynaVelo uses a loss function to reconstruct the RNA and TF motif matrices so that learned initial (latent time *t* = 0) latent states and cell-specific latent times do not greatly diverge from their specified prior distributions (**Methods**). Additional components to the loss function are used to (1) enforce the similarity between learned RNA velocities and those provided for a subset of genes by a method like scVelo and (2) ensure consistency between RNA and TF motif velocities (**Methods**).

**Figure 1.**
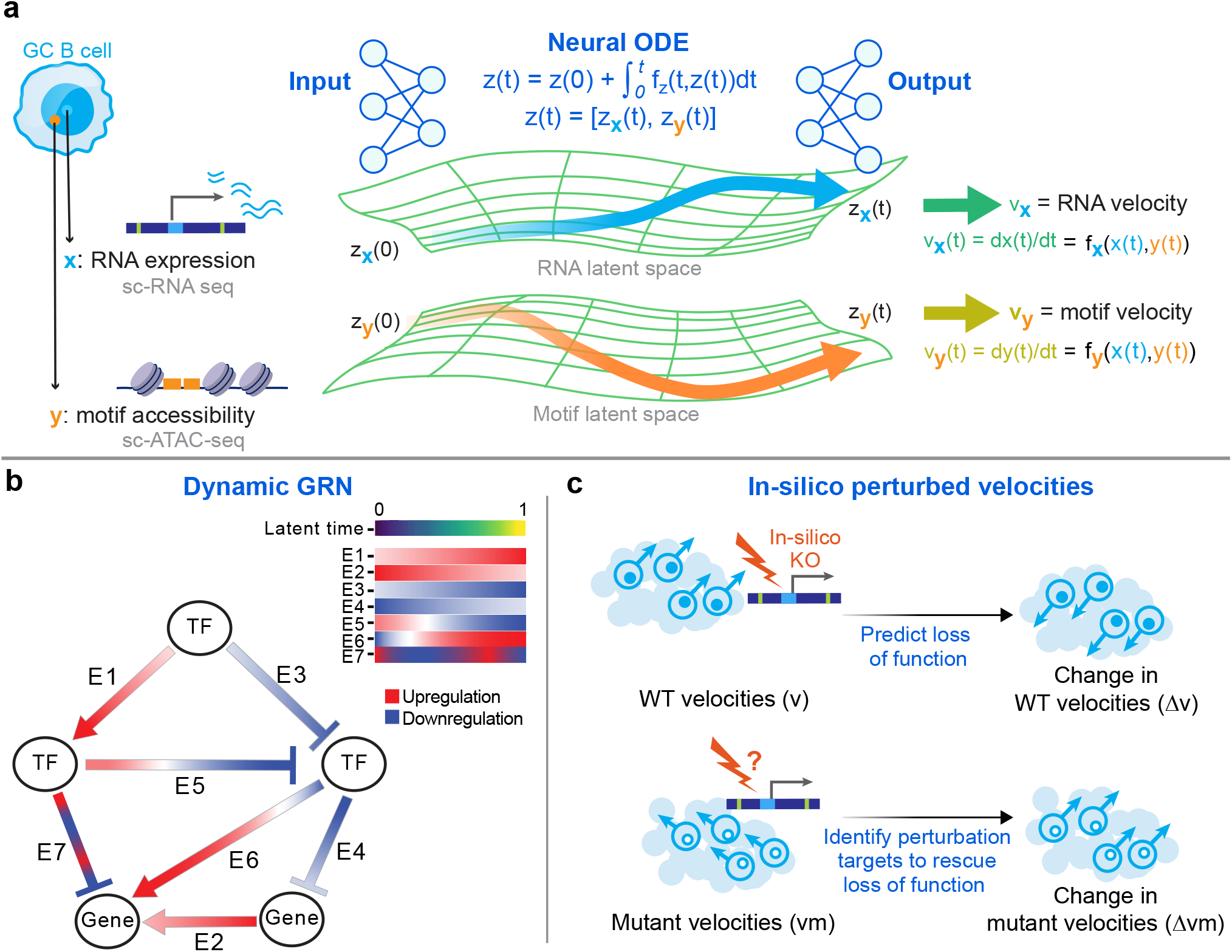
DynaVelo trains a latent neural ODE model to learn the dynamics of single cells using single-cell multiomic data. **a**. The cell-by-gene gene expression and cell-by-TF motif accessibility matrices are first embedded into a lower dimensional latent space by encoders. The initial latent state *z*(0) and latent time *t* of each cell are predicted in the latent space by neural networks and passed to a neural ODE model, which solves the ODE from time *t*=0 to time *t* and starting from *z*(0). After receiving the latent state *z*(*t*) and velocity *v*_*x*_(*t*), which forms the latent vector field, two decoders map the RNA and motif vector fields to the original RNA and motif space, reconstructing the RNA expression and motif accessibility and predicting the RNA and motif velocities. **b**. DynaVelo can learn dynamic GRNs using two approaches: Jacobian matrices and *in silico* perturbations. Dynamic GRNs, unlike static ones, capture cell-state-specific gene regulatory relationship by measuring velocity changes after RNA expression or motif accessibility changes. Jacobians measure derivatives of changes while *in silico* perturbations measure finite changes. **c**. *In silico* perturbation of wildtype samples is used to validate the models by comparing the change in velocities with the velocities of knockout (KO) samples. Similarly, *in silico* perturbation of KO samples is used to identify candidate perturbation targets to rescue loss of function mutations.

Once DynaVelo is trained (**Methods**), the model can be used for multiple downstream tasks. One key application of DynaVelo is learning dynamic and cell-state-specific GRNs. To learn dynamic GRNs, we follow two approaches. The first approach is to estimate how a small change in the expression of one gene locally changes the RNA velocity of another gene through the computation of Jacobian matrices at individual cells (**Methods**). The second GRN inference approach is through *in silico* perturbation (knockout) of RNA expression values and TF motif accessibility scores and calculation of the resulting change in the RNA and motif velocities. When perturbing TFs, we set both the TF’s gene expression and motif accessibility to their minimum possible values across cells, while for non-TF genes, we only set the RNA expression to the minimum value (**Fig. 1b**). Both approaches can yield dynamic GRNs by capturing which individual gene perturbations can lead to changes in RNA velocities of all other genes, conditioned on a specific cell state. Additionally, we can also study the effects of gene perturbations on the TF motif velocities, for example to reveal how epigenetic regulators alter the motif accessibility landscapes across cells (**Fig. 1b**).

Since DynaVelo provides a functional form of the vector field for the learned cell dynamics, the model can be used with a wildtype single-cell dataset to predict the global effect of (germline or somatic) loss-of-function alterations in disease, or given cells bearing such mutations, identify the best perturbation targets to rescue the loss-of-function phenotypes and restore homeostatic dynamics (**Fig. 1c**). For this second task, we perform *in silico* perturbation of all genes in the mutant system and see which ones change the RNA and motif velocities in a direction similar to the velocities of wildtype cells (**Fig. 1c**).

### DynaVelo learns the complex dynamics of germinal center B cells and their latent times

We first applied DynaVelo to a single-cell multiome data set profiling wildtype murine germinal center B cells (*N* = 7166 cells). To guide the dynamics, we used RNA velocity estimates from scVelo for the subset of genes with sufficient coverage, called velocity genes. DynaVelo then learned RNA velocities of all genes as well as TF motif velocities, which were visualized by projection to the corresponding RNA expression and TF motif accessibility UMAP spaces (**Fig. 2a**). The clear branching of these vector fields into two types of trajectories recapitulated the main well-known dynamics of GC B cells. In one branch, the highly proliferative centroblasts (CB, shown in green) in the dark zone of GCs progress through the cell cycle. In the second main branch, CB cells transition into the centrocytes (CC, shown in blue) in the light zone of the GC. A third important and more difficult trajectory to capture is the recirculation of some CCs from the light zone back into the dark zone, visible as flow lines that pass through the recycling CC population (CC Rec, shown in orange). These recycling CC cells have been presented with antigen by follicular dendritic cells and have received survival signals from T follicular helper (TFH) cells, then migrate back to the dark zone to undergo somatic hypermutation and improve their BCR affinity^27–29^. Finally, some CCs exit the germinal center reaction towards prememory B cells (shown in purple) or plasmablasts (shown in cyan, **Fig. 2a**), which will eventually differentiate into memory B cells and plasma cells. **Figure 2b** summarizes the major GC dynamics and trajectories together with cell type annotations.

**Figure 2.**
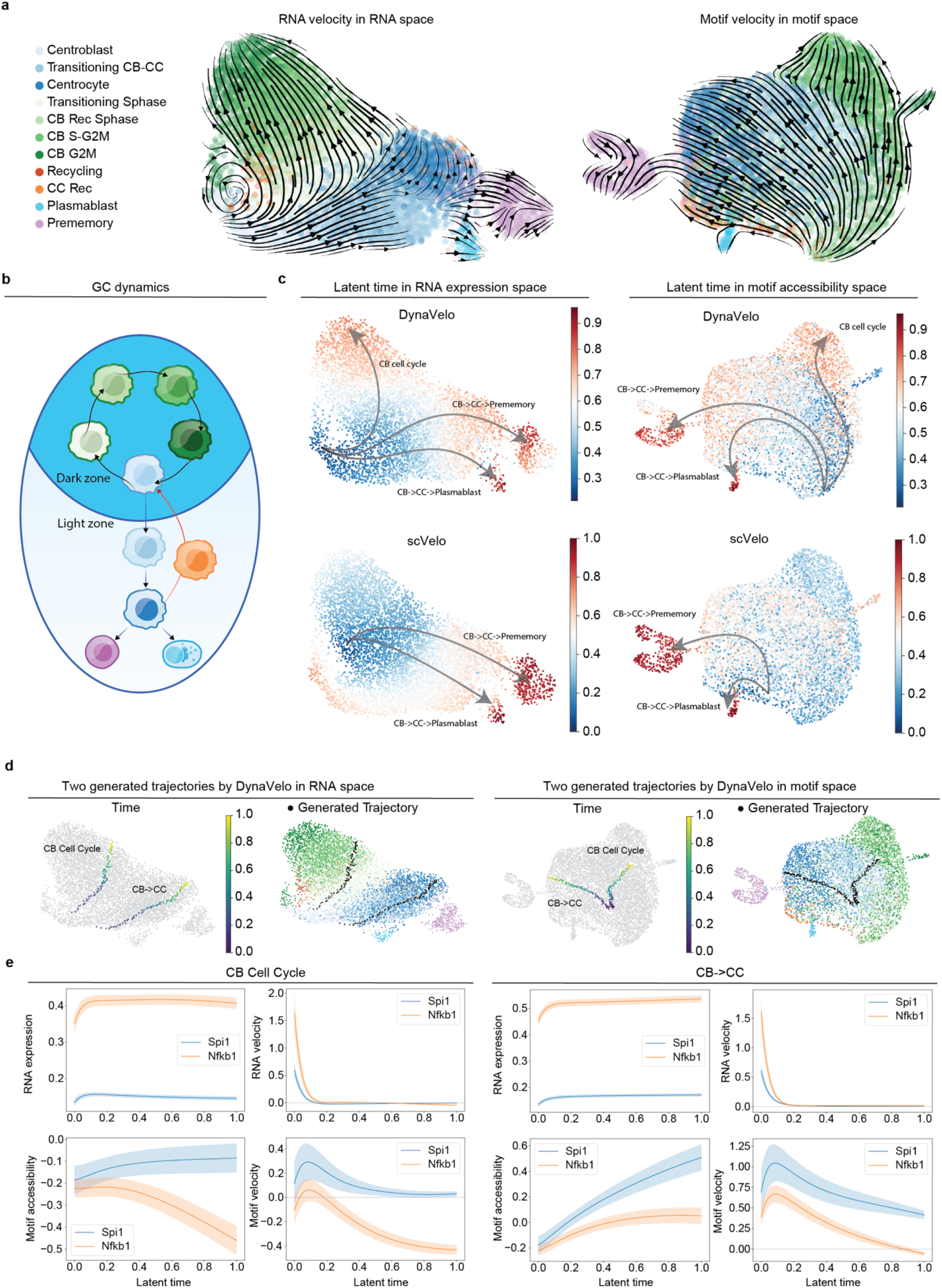
DynaVelo learns the complex dynamics of germinal center B cells and their latent times. **a**. RNA and motif velocities projected to the UMAP embeddings of RNA expression and motif accessibility, respectively, showing RNA and motif vector fields. The most important cell state transitions are uncovered: CB to CC, CB cell cycle, CC to prememory, CC to plasmablast, CC to CB migration. **b**. The summarized dynamics of GC B cells. **c**. Latent times of the cells learned by DynaVelo and scVelo shown on both RNA and motif UMAPs. DynaVelo latent times better match the dynamics of GC B cells and identify more biologically reasonable initial cells (i.e. cells with latent time *t* close to 0). **d**. DynaVelo is used to simulate trajectories capturing the dynamics of two main types: CB cell cycle and CB→CC transition. The generated RNA and motif trajectories along these simulated cells are embedded along with the input cells into the RNA and motif UMAP spaces. **e**. Dynamics of two critical GC TFs, Spi1 and Nfkb1 in two simulated trajectories, cell cycle and CB→CC. Solid lines and shades show the mean and standard errors based on 20 generated trajectories. Trends of four modalities are shown for each TF: RNA expression, RNA velocity, motif accessibility, and motif velocity. Velocity curves are the time-derivatives of the state curves.

We next plotted the latent times of cells learned by DynaVelo, shown both in RNA expression and motif accessibility UMAP spaces, and compared with scVelo’s latent times (**Fig. 2c)**. This side-by-side comparison showed DynaVelo’s latent time better recapitulated GC dynamics than those of scVelo. While both methods identified prememory B cells and plasmablasts as the terminal states, scVelo had difficulties in capturing the latent times of CBs and CCs, as they have highly dynamic transcriptional and chromatin changes, and failed to identify the major bifurcation between cells that progress through cell cycle in the dark zone and those that transition to the light zone (**Fig. 2c**). This discrepancy was even more evident in the motif space, where DynaVelo’s latent times provided a meaningful pseudo-ordering while scVelo’s latent times did not (**Fig. 2c**). Taken together, these results show that by combining RNA expression and TF motif accessibility spaces, DynaVelo’s latent times outperformed scVelo by better capturing intricate and fast-changing cell-type dynamics within GCs.

We also compared DynaVelo’s RNA velocities and latent times with MultiVelo, a velocity method that also uses both RNA expression and chromatin accessibility (**Supplementary Fig. S2**). We observed that MultiVelo’s RNA velocities did not capture the dominant transition of CBs to CCs and instead had all trajectories moving from CC to CB (**Supplementary Fig. S2a**). Moreover, unlike DynaVelo, MultiVelo failed to identify the plasmablasts as one of the GC exits and instead incorrectly identified them as a starting population (**Supplementary Fig. S2a**). This mistake was also evident in the latent times estimated by MultiVelo as they highlighted plasmablasts as the starting point, while DynaVelo’s latent times correctly identified the CBs as the starting point and plasmablasts and prememory B cells as the GC terminal states (**Supplementary Fig. S2b)**. More details on robustness and method comparison are provided in the **Supplementary Note**.

Since DynaVelo is a generative model, it can be used to generate trajectories starting from or ending at any cell and to generate cells *de novo* along these trajectories. These generated trajectories can be used to analyze cell state transitions along latent time and determine possible cell fates given a starting cell state. We used DynaVelo to generate two example trajectories, both starting in the CBs, one progressing through the cell cycle inside the dark zone, and the other one transitioning towards CCs in the light zone (**Fig. 2d**). These two trajectories are colored by the latent time points used in the neural ODEs and are projected to both RNA expression and motif accessibility spaces. These generated latent trajectories were mapped by the decoders to the original high dimensional RNA expression and motif accessibility spaces to generate 100 cells, which were then embedded together with the observed cells to their respective UMAP spaces (**Fig. 2d**).

Trajectory generation is a unique feature of DynaVelo that can be used to understand the vector field over cells in both RNA and motif spaces and to examine gene expression and TF motif accessibility dynamics in different trajectory types. For instance, we examined the dynamics of Spi1 and Nfkb1, two critical TFs in GC B cells, along both CB→CC and CB cell cycle trajectories (**Fig. 2e**). Here the velocity (RNA and motif) curves are the time derivatives of their corresponding state (RNA expression and motif accessibility) curves. A recent study showed using functional assays that Spi1 is required for transitioning of CBs to CCs and acts as a pioneer factor to open chromatin, recruiting chromatin remodeling complexes and subsequently enabling binding of Nfkb factors^30^. Our results validated these findings by showing that Spi1 and Nfkb1 accessibilities are monotonically increasing from CB to CC, but not in CB cell cycle progression where Spi1 motif accessibility is stable while Nfkb1 is monotonically decreasing (**Fig. 2e**). Interestingly, motif accessibility demonstrated more dynamics than RNA expression for these TFs, confirming the importance of including this modality for resolving TF activity in the germinal center. These results show that DynaVelo was able to determine the temporal sequence of TF activity and binding *in silico*, with the potential to discover novel TF co-dependencies.

### DynaVelo identifies driver TFs for the main GC trajectory types

Having confirmed that DynaVelo identifies major classes of GC B trajectories – CB progression through cell cycle, CB to CC transition, and exits towards prememory B cells and plasmablasts – we next delved into each trajectory type more carefully by analyzing dynamic patterns of TF expression and motif accessibility along latent time. To this end, we first partitioned the cells based on their trajectories, sorted them according to their latent time, and plotted the RNA expression and motif accessibility temporal patterns of important GC TFs (**Fig. 3a**). We expected that driver TFs for each cell type would likely display monotonic patterns of TF expression and motif accessibility, both increasing and decreasing, within the cell type, and that the temporal order of TF activity might also play a role in cell fate decisions.

**Figure 3.**
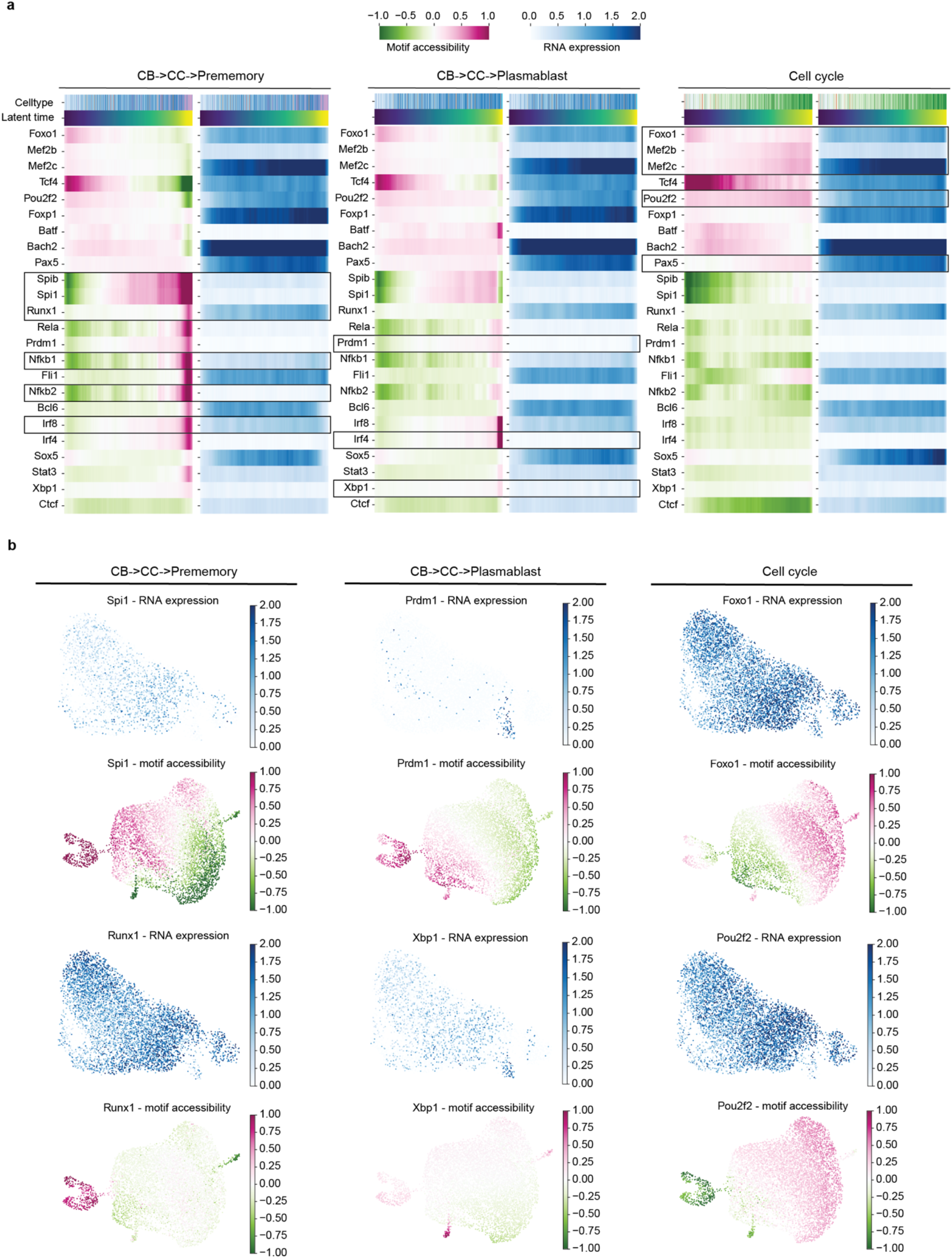
DynaVelo identifies driver TFs for key GC trajectories. **a**. The temporal trends of motif accessibility and RNA expression for essential germinal center TFs across DynaVelo latent time in three main trajectory types: CB→CC→prememory, CB→CC→plasmablast, and CB cell cycle. **b**. RNA expression and motif accessibility of several important TFs in each GC trajectory type shown respectively on RNA and motif UMAPs. Both RNA expression and motif accessibility of TFs are required to understand their roles in each trajectory.

For the prememory B cell state, in addition to Spi1/Spib and Nfkb1/2, TFs including Runx1 and Irf8 displayed an increasing pattern, suggesting their possible roles in the progression to CC to prememory (**Fig. 3a**,**b**)^30,31^. The RNA expression and motif accessibility temporal patterns of Spi1 and Runx1 are depicted on the corresponding UMAP embeddings in **Fig. 3b**, showing that the accessibility of Spi1 gradually increases from CB to CC to prememory B cells but not in plasmablasts. The RNA expression of Spi1 is also absent in plasmablasts. Notably, the Runx1 motif is not accessible in the GC but becomes accessible at a later stage when committing to the prememory B cell fate (**Fig. 3b**).

Based on similar DynaVelo analysis, some of the important TFs driving the plasmablast fate include Prdm1, Irf4, and Xbp1^31^ (**Fig. 3a**). Even though TF motif accessibility is a good proxy for single-cell TF activity, it alone appears insufficient to identify TF drivers; we found that a TF’s RNA expression was also required to accurately infer its role. For example, Prdm1 is accessible in both prememory B cells and plasmablasts but is only expressed in plasmablasts, which shows that Prdm1 is only responsible for plasmablast differentiation (**Fig. 3b**). Similarly, Xbp1 demonstrates both higher accessibility and expression in plasmablasts, consistent with the fact that Xbp1 is a plasma cell TF.

Contrary to prememory B cells and plasmablasts for which important TFs have been studied, less is known about CB-specific drivers of cell cycle in the dark zone. DynaVelo temporal patterns identified Foxo1, Mef2b, Mef2c, and Pou2f2 as TFs that potentially play a role in CB cell cycle progression (**Fig. 3a**), all of which are well-known GC TFs based on the literature^31^. These TFs are expressed in both CBs and CCs, but their motif accessibility increases with CB cell cycle progression and decreases when transitioning towards CCs (**Fig. 3a**,**b**). These observations are consistent with known biology of these TFs, which are important for maintaining the GC reaction but play a limited role in GC cell differentiation towards prememory B or plasmablast fates.

### Heterozygous mutations in Arid1a and Ctcf disrupt the normal trajectories of germinal center B cells

In order to determine if DynaVelo can capture changes in cell-fate dynamics upon perturbation of key regulators of the GC reaction, we next turned from wildtype GC B cells to those with reduced expression of *Arid1a* and *Ctcf*, which respectively encode a DNA-binding subunit of the BAF chromatin remodeling complex and the insulator protein Ctcf that acts as both a TF and mediator of 3D chromatin looping. To this end, we applied DynaVelo to two multiome samples, *Arid1aHet* (*N* = 6716 cells) and *CtcfHet (N* = 4688 cells), with heterozygous conditional deletion of *Arid1a* and *Ctcf*, respectively, in GC B cells^30^. Latent times learned for cells in these samples were similar to the wildtype samples in that they identified a subset of CBs as root cells and prememory B cells and plasmablasts as terminal cells (**Fig. 4a**). The learned RNA velocities captured some of the main GC trajectories including CB cell cycle, CB to CC transition, and differentiation towards prememory B cells and plasmablasts (**Fig. 4a**). However, one important expected trajectory was lost in both samples, namely the recirculation of CCs to CBs. This is an important physiological trajectory in the GCs, as it is necessary for B cell receptor (BCR) affinity maturation prior to differentiation to either memory B cells or plasma cells. As shown before, loss of this trajectory most likely leads to a premature exit from the GC at the prememory B cell state^30^. Indeed, the percentage of prememory B cells in the mutant samples (20.09% in *Arid1aHet*, 25.45% in *CtcfHet*, and 44.96% in *Arid1aHom*) is increased compared to wildtype (5.42%) (**Supplementary Fig. S1**). This indicates that in the mutant samples, GC B cells do not undergo multiple rounds of somatic hypermutation in order to improve their BCR affinity, and consequently they have a faster exit as premature memory B cells.

**Figure 4.**
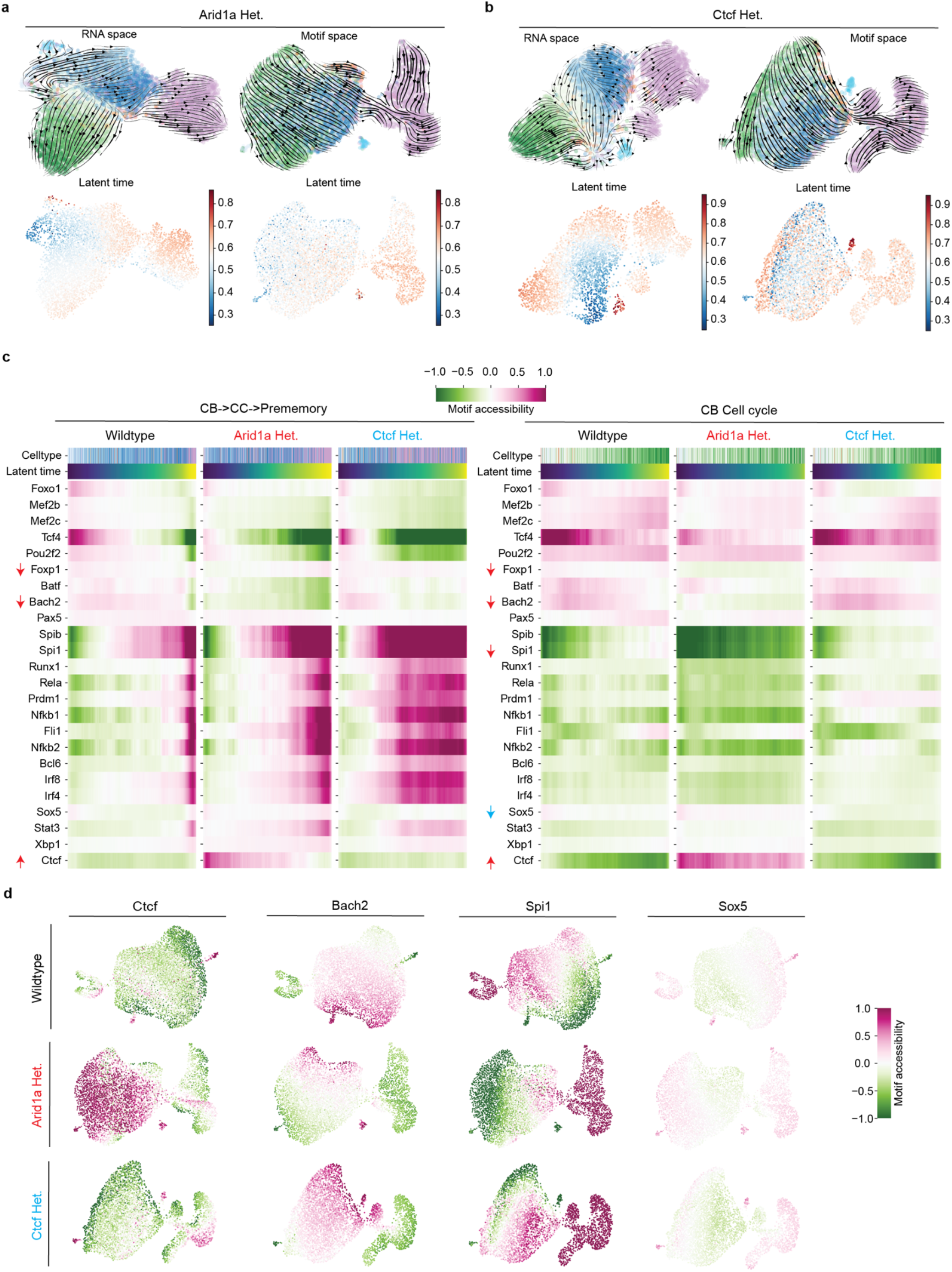
Heterozygous mutations in Arid1a and Ctcf disrupt the normal trajectories of germinal center B cells. **a**. RNA and motif velocities of *Arid1aHet* GC B cells and DynaVelo’s latent times projected to the RNA and motif UMAPs. **b**. RNA and motif velocities of *CtcfHet* GCB cells and DynaVelo’s latent times projected to the RNA and motif UMAPs. In both (**a**) and (**b**), the critical recycling of CCs from the light back to the dark zone is lost, leading to either apoptosis or to germinal center exit as premature prememory B cells. **c**. TF motif accessibility trends of wildtype, *Arid1aHet*, and *CtcfHet* GC B cells in two main trajectory types: CB→CC→prememory and CB cell cycle. The dynamics of some TFs change more than others in the mutant GCs, highlighted by red and blue arrows for *Arid1aHet* and *CtcfHet*, respectively. **d**. Motif accessibility of TFs whose temporal patterns change more in *Arid1aHet* and *CtcfHet* GCs.

To identify key transcription factors disrupted in mutant GCs, we analyzed their altered dynamics in CB cell cycle progression and the CB→CC→prememory trajectory type compared to WT (**Fig. 4c**). While many TFs displayed subtle changes in dynamics, several TFs were especially notable in their strong changes in motif accessibility in the *Arid1aHet* sample, including Foxp1, Bach2, Spi1, and Ctcf (**Fig. 4c**). The accessibility of Foxp1 and Bach2 decreases in *Arid1aHet* in both trajectory types compared to WT (**Fig. 4c,d**), while the accessibility of Spi1 decreases in *Arid1aHet* during CB cell cycle progression (**Fig. 4c,d**). Interestingly, the accessibility of Ctcf increases significantly in both CBs and CCs in *Arid1aHet* compared to WT (**Fig. 4c,d**). This observation implies that Arid1a and Ctcf may function antagonistically within the germinal center. Overall, the *CtcfHet* displayed a stronger phenotype than *Arid1aHet*, in that TFs exhibited a stronger change in TF motif activity relative to WT. Additionally, we highlighted Sox5 as a potential TF whose accessibility decreases slightly in *CtcfHet* compared to WT, especially during CB cell cycle (**Fig. 4c,d**).

### DynaVelo uncovers dynamic TF relationships in GCs upon loss of chromatin regulators Arid1a and Ctcf

So far, we have shown that DynaVelo is able to learn the main trajectories of GCs and their potential driver TFs and to identify the disrupted trajectories of mutant GC B cells. We next used DynaVelo to derive dynamic GRNs through examination of Jacobian matrices at cells of each trajectory type or through *in silico* perturbation of genes to understand how RNA expression and TF motif accessibility regulate the RNA and motif velocities. In particular, we used this analysis to identify dynamic TF-target gene regulatory relationships along trajectories. We restricted to showing known gene regulatory relationships from literature^31^ that are recovered based on the Jacobian matrix *J*_*xx*_, which measures how the change in RNA velocity of one gene depends on the change in expression of another gene, if TF motif accessibilities are left unchanged (**Fig. 5a**). Positive and negative Jacobian values correspond to upregulation and downregulation, respectively. One limitation of this assessment is that known TF-to-gene regulatory relationships, i.e. GRN ‘edges’, from the germinal center literature^31^ are reported as static rather than cell-state-dependent and dynamic. DynaVelo is able to extract the dynamics of gene regulatory relationships at the single-cell level and display temporal patterns along latent time in CB cell cycle and CB→CC→prememory trajectories in WT, *Arid1aHet*, and *CtcfHet* samples (**Fig. 5a,b**). One of the important regulatory edges in the GC is Spi1→Bcl6 upregulation^31^, which matches the DynaVelo Jacobian in the WT CB→CC→prememory trajectory type. Notably, this regulatory relationship is lost in both *Arid1aHet* and *CtcfHet* in this trajectory type (**Fig. 5a**).

**Figure 5.**
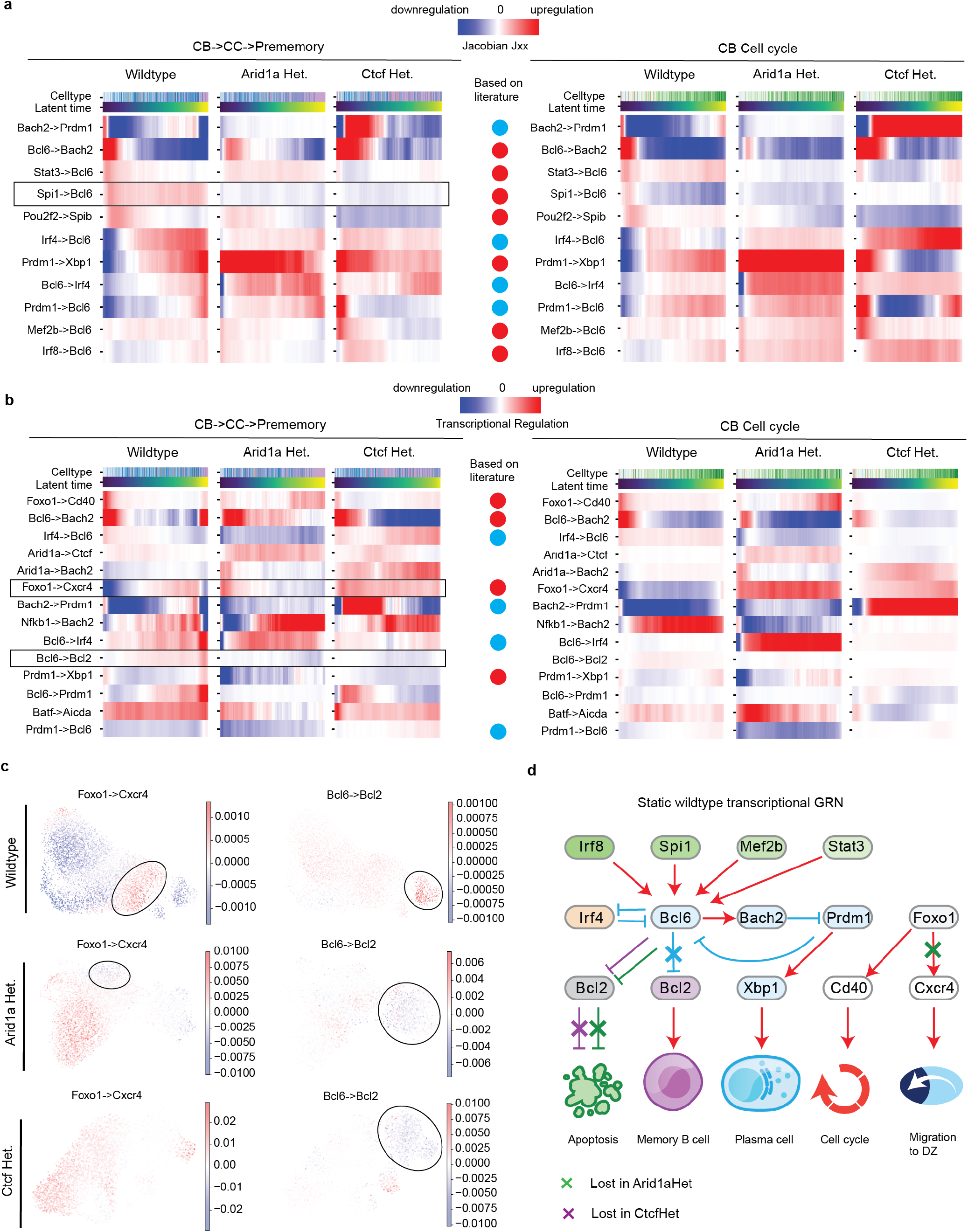
DynaVelo learns dynamic GRNs of wildtype GCs and identifies disrupted gene regulatory relationship in Arid1a and Ctcf mutant GC. **a**. Dynamic GRN edges for wildtype, *Arid1aHet*, and *CtcfHet* GCs in two trajectory types (CB→CC→prememory and CB cell cycle) based on analysis of the Jacobian Jxx. The majority of the wildtype dynamic regulatory edges match the known static GC GRN edges but provide more information about cell state in which they hold. The wildtype edge, Spi1→Bcl6 upregulation, is missing both in *Arid1aHet* and *CtcfHet* GCs. **b**. Dynamic GRN edges for wildtype, *Arid1aHet*, and *CtcfHet* GCs in two trajectory types (CB→CC→prememory and CB cell cycle) by *in silico* perturbations. Two important edges are lost in mutant GCs: Foxo1→Cxcr4 upregulation is lost in CCs in *Arid1aHet*, which is required for the migration of CCs from the light to the dark zone; Bcl6→Bcl2 downregulation is gained in prememory B cells in *Arid1aHet*, preventing normal differentiation toward memory B cells and priming the cells for apoptosis. **c**. The regulatory edges Foxo1→Cxcr4 and Bcl6→Bcl2 shown in the RNA UMAPs of wildtype, *Arid1aHet*, and *CtcfHet* GCs. **d**. A minimal static GRN of the wildtype GC is summarized from the dynamic GRN analysis. Red arrows represent upregulation; blue arrows, downregulation. The Bcl6→Bcl2 repressive edge is lost in the WT GC to enable differentiation to memory B cells (shown in blue cross). The edges that are lost or gained in mutant GC B cells are shown in purple and green lines and crosses.

A second approach to investigate TF regulatory effects is through *in silico* perturbation, namely setting both the TF RNA expression and motif accessibility to their minimal values and measuring the change in RNA velocities for all other genes. Note that *in silico* perturbation analysis captures slightly different regulation patterns. In calculating *J*_*xx*_, only TF RNA expression is perturbed by adding a small value (*ϵ* = 10^−4^), while motif accessibility is unchanged. For *in silico* perturbations, both TF RNA expression and motif accessibility are perturbed by setting them to their minimum values across cells. **Figure 5b** shows several important and known gene regulatory relationships in GCs^31^, along with the cell states/trajectory types in which they hold. Similar to the Jacobian analysis, *in silico* perturbation identified regulatory edges that are lost in *Arid1aHet* and *CtcfHet* mutant GCs. One critical transcriptional GRN edge in wildtype GCs is Foxo1→Cxcr4 upregulation, which is necessary for the maintenance of the GC and migration of CCs to the dark zone^31^. Notably, Foxo1→Cxcr4 upregulation is lost in CCs in *Arid1aHet* but instead is observed in CB cell cycle, as compared to WT (**Fig. 5b,c**). This lost regulatory edge may be a reason why CCs in *Arid1aHet* lose their capability to migrate back to the dark zone and instead exit as premature prememory B cells, as described in our earlier analysis and experimentally validated^30^.

Interestingly, the inferred sign of Bcl6→Bcl2 regulatory influence in the transition of CCs to prememory B cells is different in both mutants compared to wildtype (**Figure 5b,c**). Bcl6 is a master transcriptional repressor that downregulates expression of the anti-apoptotic TF Bcl2^31^. This repression is relieved in CCs that receive T cell help, allowing increased expression of Bcl2 to promote B cell survival. In wildtype, the sign of the Bcl6→Bcl2 becomes positive, opposite of what we observe in both *Arid1aHet* and *CtcfHet*. suggesting that ARID1A and CTCF regulate GC dynamics through control of Bcl6-Bcl2 axis.

To summarize our findings, we assembled a minimal GRN governing the transcriptional regulation of critical GC TFs and highlighted regulatory relationships that are compromised in *Arid1aHet* and *CtcfHet* GCs (**Fig. 5d**).

### In silico perturbations of Arid1a mutant GCs identify candidate targets to rescue the loss of function

One powerful application of DynaVelo is the ability to perform *in silico* loss-of-function perturbations and determine their effects on GC dynamics and phenotype. We next aimed to assess how accurately we could predict the impact of genetic loss-of-function on RNA and motif velocities, and also to determine which additional loss-of-function perturbations would rescue the phenotypes of *Arid1a* loss. Notably, the change in the RNA and motif velocities (delta velocities) after *in silico* perturbation of *Arid1a* in the WT DynaVelo model suggests a decreased connectivity of CB and CCs potentially reflecting the loss of the CC-to-CB recycling trajectory. Instead the aforementioned trajectory between CCs and prememory B cells became highly prominent (**Fig. 6a**), suggesting a preferential prememory B cell differentiation and subsequent premature GC exit.

**Figure 6.**
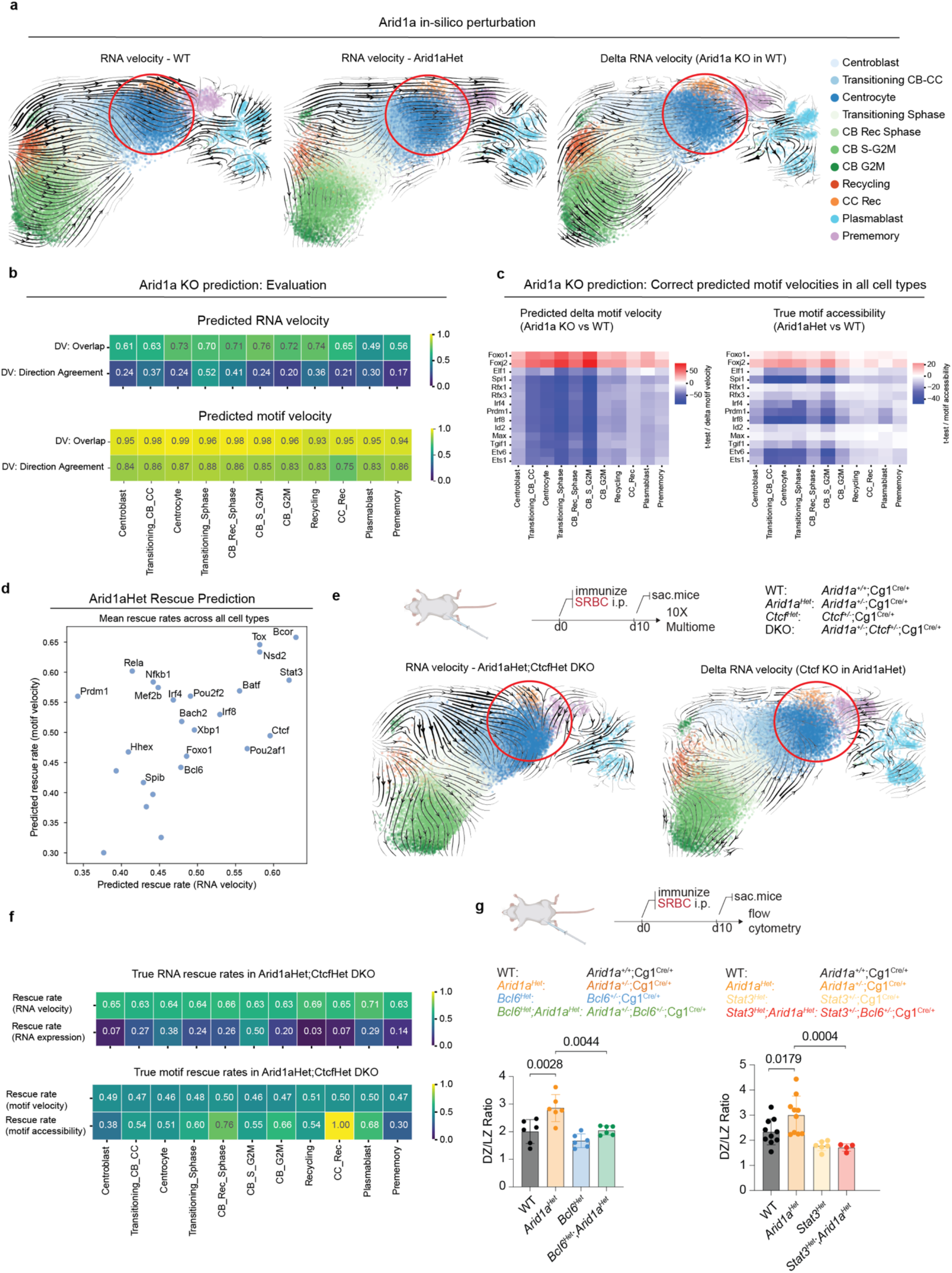
In silico perturbations of Arid1a mutant GCs identify candidate targets for rescuing the loss of function. **a**. RNA velocities of WT, *Arid1aHet*, and delta RNA velocities after *in silico* perturbation of *Arid1a* in the WT cells. Delta velocities show the difference between the velocity vectors before and after perturbation. These delta velocities are projected to the RNA UMAP derived from all samples, where the arrows indicate the direction toward which the cells are inclined to move after the perturbation. These trajectories show that the cells tend to transition from CB to CC and CC to CB recirculation is lost. **b**. Evaluation of predicted Arid1a KO in WT cells. Differential velocity (DV) overlap and direction agreement show how much the significant predicted RNA and motif velocities overlap with the true ones and how much they agree in sign (direction). **c**. The complete list of TFs for which *Arid1a* delta motif velocities match the motif velocity and accessibility changes in all cell types when comparing wildtype and *Arid1aHet*. **d**. Predicted rescue rates of RNA and motif velocities after *in silico* perturbation of essential TFs in *Arid1aHet*. These rates are the mean of rescue rates across all cell types. The TFs at the top right are the best perturbation targets for rescuing the loss of function phenotype in *Arid1aHet*. **e**. RNA velocities of *Arid1aHet*;*CtcfHet* DKO cells and delta RNA velocity after *in silico* perturbation of Ctcf in the *Arid1aHet* cells. **f**. True rescue rates of RNA and motif velocities, RNA expression, and motif accessibility in the DKO GC in each cell type. **g**. Bcl6 and Stat3 heterozygous KO in *Arid1aHet* GC (leading to two DKO GCs: *Arid1aHet*;*Bcl6Het* and *Arid1aHet*;*Stat3Het*) rescue DZ/LZ ratio phenotype of *Arid1aHet* GCs, which is higher than WT GCs. Overall 54 samples are used. The number of samples in Bcl6 experiment: WT: 6, *Arid1aHet*: 6, *Bcl6Het*: 6, *Arid1aHet*;*Bcl6Het* DKO: 6. The number of samples in Stat3 experiment: WT: 10, *Arid1aHet*: 10, *Stat3Het*: 6, *Arid1aHet;Stat3Het* DKO: 4.

For a more quantitative assessment, after *in silico* perturbation of *Arid1a*, we first performed a statistical test to identify RNA and motif delta velocities in each cell type whose mean was significantly different from zero, either positive or negative, corresponding to significant velocity increases and decreases, respectively (**Methods**). Then we tested if the mean velocities were significantly difference between *Arid1aHet* and WT GCs in each cell type (**Methods**). We defined two metrics to assess how the predicted and true significant (*p*_*adj*_ < 0.05) differential velocities (DV) match: DV overlap and DV direction agreement. DV overlap is defined as the Jaccard index (size of intersection divided by the size of union) of predicted and true DV sets. DV direction agreement is defined as the fraction of the velocities with the same sign (positive or negative) in the intersected DV set. **Figure 6b** shows the results of *Arid1a* KO in the WT GCs, where we see that the predictions of motif velocities are more accurate by these metrics than the RNA velocities in all cell types. **Figure 6c** shows the correctly predicted TF motifs for which the predicted and true velocity changes match each other as well as the motif accessibility changes in all the cell types. Of note is the important GC TFs such as upregulation of Foxo1 (mostly known as dark zone TF) and downregulation of Spi1, Irf4 and Prdm1.

As a final application of DynaVelo, we performed *in silico* perturbation of important GC TFs and epigenetic regulators in the mutant GC models and identified which perturbation targets could potentially rescue the loss of function in *Arid1aHet* GC B cells (**Fig. 6d)**. A velocity was counted as rescued in a given cell type if the *in silico* perturbation significantly shifted it towards its wildtype value. For each candidate perturbation in the *Arid1aHet* model, we computed the number of rescued RNA and motif velocities across cell types and defined the rescue rate as the DV direction agreement, and **Fig. 6d** shows the mean rescue rates (RNA and motif) for all cell types. This approach identified the perturbation of TFs like *Tox, Bcor, Ctcf, Stat3* and *Bcl6* as potentially leading to rescue of the *Arid1aHet* phenotype (**Fig. 6d**). Notably, this *in silico* prediction of *Ctcf* as a rescue perturbation suggested antagonizing effects of *Ctcf* and *Arid1a* in GC biology. Importantly, recent work has established a mechanistic basis for this predicted antagonism and demonstrated a phenotypic rescue of *Arid1aHet* DZ increase in *Arid1aHet;CtcfHet* double heterozygous knockout (DKO) animals (by flow cytometry; Debek, Meydan et al., unpublished manuscript, **Related Manuscript File**), confirming a key result from our DynaVelo rescue analysis.

To validate this rescue at the gene expression and chromatin level, we performed single-cell multiome on GC cells from *Arid1aHet;CtcfHet* DKO mice as well as control animals. In the *Arid1aHet;CtcfHet* DKO GCs, the DynaVelo-inferred RNA velocity field showed that the rapid CC to prememory exit characteristic of *Arid1aHet* was hindered and partially reversed (**Fig. 6e**, left panel). This experimental outcome closely mirrored the delta RNA velocity predicted by *in silico Ctcf* perturbation of *Arid1aHet* cells (**Fig. 6e**, right panel), confirming that DynaVelo’s prediction recapitulated the dynamics of the genetic double mutant. To quantify rescue across cell types, we computed the fraction of features that shifted from *Arid1aHet* toward wildtype values in the DKO across four modalities: RNA expression, RNA velocity, motif accessibility and motif velocity (**Fig. 6f, Methods**). We refer to this fraction as the true rescue rate, in contrast to the predicted rescue rates from *in silico* perturbation reported above. Rescue was observed across all cell types but varied in magnitude. In CCs, the cell type whose dynamics are most disrupted in *Arid1aHet*, the rescue rates were 38% (RNA expression), 64% (RNA velocity), 51% (motif accessibility) and 46% (motif velocity).

A defining feature of *Arid1aHet* GCs is an increased dark zone to light zone (DZ/LZ) ratio, consistent with the premature exit of light zone cells toward the prememory fate described above (**Fig. 6g**). While DynaVelo *in silico* perturbation analysis nominated *Bcl6* and *Stat3*, alongside *Ctcf*, as candidate rescue targets for *Arid1a* loss, an independent *in vivo* test is required to establish that these predictions reflect functional genetic interactions. We hypothesized that if DynaVelo predictions are correct, compound heterozygous loss of either *Bcl6* or *Stat3* with *Arid1a* should restore the DZ/LZ ratio toward wildtype. To test this, we generated two entirely new mouse models carrying double heterozygous mutations, *Arid1aHet;Bcl6Het* and *Arid1aHet;Stat3Het*, and performed immunophenotyping of GC B cells from each strain. The DZ/LZ ratio was restored to wildtype levels in both compound heterozygous backgrounds (**Fig. 6g**), directly confirming the rescue predicted by DynaVelo.

Together with the independent rescue of the same phenotype by *Ctcf* heterozygous loss in *Arid1aHet* mice (Debek, Meydan et al., unpublished manuscript), these experiments establish that all three rescue targets nominated by DynaVelo (*Ctcf, Bcl6*, and *Stat3*) functionally rescue an *in vivo* phenotype of *Arid1a* loss. These results demonstrate that DynaVelo’s *in silico* perturbation framework can identify genetic interactions and nominate experimentally tractable rescue strategies for loss-of-function disease alleles, reducing the need for animal models.

## Discussion

A fundamental goal in biology is to understand the dynamic GRNs that govern cell state transitions in evolving cellular systems and to predict how cell-fate decisions change when the network is perturbed. Since most single-cell genomics technologies are destructive, cell dynamics and GRNs are typically inferred from data at a single snapshot of time, using pseudotime strategies such as those based on RNA velocity estimates. However, these velocity estimates are prone to errors and can generally be computed only for a relatively small subset of genes. Another key challenge is to distinguish causal drivers from non-causal ones, as single-cell data provides a high dimensional (and sparse) space of observables comprising many confounding factors. With the advent of single-cell multiome technology, it is now possible to exploit single-cell TF motif activities derived from scATAC-seq in addition to gene expression and RNA velocity in dynamic models of evolving cellular systems. We developed DynaVelo as a robust generative deep learning model that uses single-cell chromatin accessibility and transcriptomic readouts at a single time point to infer dynamic gene regulation patterns, and we applied this model to dissect the complex *in vivo* dynamics of WT and mutant GC B cells.

DynaVelo uses a neural ODE framework where RNA expression and TF motif accessibility inputs are each mapped to latent spaces in which the cell dynamics are learned. Partial RNA velocities from a tool like scVelo are supplied as input to provide pseudotemporal guidance, and the model learns RNA velocities for all modeled genes as well as TF motif velocities. Through Jacobian analysis and *in silico* perturbation of TFs, DynaVelo can reveal *dynamic* GRNs, where single-cell regulatory edges between TFs and target genes depend on cell state, moving beyond traditional GRN inference methods that present a single static network per dataset or cell type. Moreover, DynaVelo provides a powerful generative deep learning framework for modeling and intervening in dynamic cellular systems, enabling the generation of cellular trajectories and the analysis of *in silico* genetic perturbations at both the expression and motif accessibility levels.

Other recent work has used machine learning models on single-cell data to infer continuous vector fields and model cell dynamics. For example, Dynamo^20^ infers absolute RNA velocity, predicts cell fates and perturbation outcomes, and extracts regulatory relationships using differential geometry. However, training this model requires single-cell RNA-seq paired with metabolic labeling which is not feasible in many experimental systems. Another recent model scDiffEq^21^ uses drift-diffusion modeling of cell dynamics but only uses RNA expression without chromatin accessibility information and thus fails to leverage scATAC-derived TF activity dynamics. By contrast, DynaVelo exploits single-cell multiome to powerfully model how gene expression and chromatin accessibility, as summarized by TF motif activities, influence each other in cell dynamics, enabling the accurate prediction of loss-of-function and rescue perturbations. Another multiome-based model MultiVelo also uses both modalities, scRNA-seq and scATAC, to model the RNA velocities of cells, which extends the gene-based ODE models of scVelo (spliced/unspliced counts) to incorporate a third ODE for gene-level chromatin accessibility. However, MultiVelo (like scVelo) does not provide a functional form for the underlying vector field — i.e. it allows us to draw a picture of trajectories, but there is no way to intervene in the system to ask how various perturbations would change the dynamics/trajectories. By contrast, DynaVelo learns a functional form of cell dynamics (instantiated by a neural ODE model), allowing us to extract dynamic gene regulatory networks, to predict the impact of loss-of-function perturbations, and to predict how to rescue loss-of-function phenotypes.

We first used DynaVelo in the WT GCs to uncover main dynamics and trajectories of the GC reaction. GC B cells experience rapid and extensive shifts in their chromatin and transcriptional programs as they cycle back and forth between rounds of proliferative bursting in the GC dark zone and selection in the light zone and exit the GC reaction as memory B cells or plasma cells^32^. These rapid cell-fate shifts are dictated by constantly evolving chromatin patterns, presenting a challenge to existing trajectory and velocity inference algorithms. DynaVelo recovered expected dynamics of GCs and the driver TFs of CB cell cycle progression, CB to CC transition, and prememory, and plasmablast exits. We showed that Spi1 and Nfkb1 are two driving TFs required for CBs to transition to CCs and then to prememory B cells, consistent with previous findings^30^. Next, we applied DynaVelo to mutant GCs and identified that the main lost transition in both *Arid1aHet* and *CtcfHet* GC dynamics is the migration from the light zone back to the dark zone, a critical step required for B cell affinity maturation before differentiation to prememory B cells or plasmablasts. As a result, CCs either undergo apoptosis (dead-end trajectories) or transition towards premature prememory B cells. This is in accordance with the observation that the percentage of prememory B cells is higher in the mutant GCs and that *Arid1a* haploinsufficiency in the GC leads to a fast exit as premature prememory B cells^30^. Through analysis of TF motif accessibility along inferred latent time, we identified TFs whose dynamics were disrupted in mutant GCs, determining that Bach2, Foxp1, and Spi1 become less accessible and Ctcf becomes highly accessible in *Arid1aHet*.

Beyond resolving latent time ordering of cells, DynaVelo learns a functional form of the vector field underlying cell dynamics, enabling the inference of dynamic GRNs in the GC through Jacobian and *in silico* perturbation analysis. We showed that DynaVelo recovered TF-gene regulatory relationships in the WT GC that matched known biology and identified which regulatory edges are lost or gained in mutant GCs. We found that two important TF-gene edges are lost in the mutant setting. First, upregulation of Cxcr4 by Foxo1 is essential for migration of CCs in the light zone back to the dark zone, and this regulatory edge is lost in *Arid1aHet* CCs, leading to fast and premature exit from the GC. Second, Bcl6 repression of Bcl2 is relieved in CCs receiving T cell help prior to exit towards prememory B cells^31^ in wildtype GCs, but this negative Bcl6 to Bcl2 regulatory relationship is apparently retained in both mutants. Finally, we used *in silico* perturbations in the mutant GCs to identify candidate perturbation targets potentially leading to rescue of the loss of function in *Arid1aHet*, with *Tox, Bcor, Ctcf, Stat3* and *Bcl6* as potential candidates. We then experimentally validated that heterozygous loss of *Ctcf, Stat3*, and *Bcl6* indeed rescued the dark zone phenotype in *Arid1aHet*. DynaVelo thus provides a powerful generative neural ODE framework for modeling cell dynamics from single-cell multiome data, enabling identification of TF drivers of cell state transitions, inference of dynamic GRNs, and prediction of loss-of-function and rescue genetic perturbations.

## Methods

### Mice

All animal work was carried out in accordance with the Guide for the Care and Use of Laboratory Animals and the policies of Weill Cornell Medicine, with oversight consistent with AAALAC International accreditation standards. Protocols were reviewed and approved by the Research Animal Resource Center and the Institutional Animal Care and Use Committees of Weill Cornell Medicine and Cornell University (protocols #2011-0031 and #2025-0008; PI: Barisic). Animals were housed in a specific pathogen-free vivarium on a 12 h light/dark cycle, with temperature held at 20–22°C and relative humidity at 40–60%, and were provided food and water ad libitum. Unless stated otherwise, experiments used age- and sex-matched mice between 8 and 16 weeks of age, with both males and females represented in every experimental group; no sex-dependent effects were detected. Arid1a (027717), Cg1Cre (010611), and Stat3 (031875) lines were obtained from The Jackson Laboratory. The *CtcfHet* (Ctcf^+/−^;Cgamma1^Cre/+^) line was generously provided by the laboratory of Dr. Gregoire Lauvau (Albert Einstein College of Medicine). Bcl6 mice (Bcl6^+/−^; Cgamma1^Cre/+^) were generated and published by Dr. Ari Melnick^33^. Prior to use, the Arid1a mouse line was backcrossed to C57BL/6J for a minimum of six generations as Jackson Laboratory strain was in a mixed genetic background. The Cgamma1-Cre allele was used in heterozygous form throughout.

### Immunizations

Mice received an intraperitoneal (i.p.) injection of 500 μL of a 2% suspension of sheep red blood cells in sterile 1× DPBS, freshly prepared from sheep blood in Alsever’s solution (Cocalico Biologicals, 20-1334A).

### Flow cytometry

Harvested spleens were mechanically dissociated and the resulting cell suspensions passed through a 40 μm strainer. Mononuclear cells were then enriched by density gradient centrifugation over Ficoll-Paque Premium (Cytiva, 17-5442-02). Single-cell suspensions were resuspended in PBE (PBS supplemented with EDTA) containing rat anti-mouse CD16/CD32 (clone 2.4G2, BD) at 0.5 μg/mL and incubated on ice for at least 5 min to block Fc receptors. Samples were stained on ice for 20 min with fluorochrome-conjugated anti-mouse antibody cocktails. Antibody mixes were prepared in PBE, except when the cocktail contained two or more BV- or BUV-conjugated reagents, in which case staining buffer consisted of a 50:50 mixture of Brilliant Stain Buffer (BD, 566349) and PBS containing 0.5% BSA (no EDTA), in order to avoid polymer dye interactions. Stained cells were washed twice, and DAPI (ThermoFisher, D1306) was added immediately before acquisition at a final concentration of 0.5 μg/mL for dead-cell exclusion. Samples were run on a BD Symphony A3 cytometer, and data were processed in FlowJo. Antibodies and dilutions used: B220-APC (RA3-6B2, BD 553092, 1:100); B220-BV786 (RA3-6B2, BD 563894, 1:300); CD138-BUV737 (281-2, BD 564430, 1:500); CCR6-PerCP-Cy5.5 (29-2L17, BioLegend 129809, 1:300); CD11b-APC (M1/70, BioLegend 101212, 1:400); CD21-APC (7E9, BioLegend 123412, 1:500); CD21-PECy7 (7E9, BioLegend 123420, 1:500); CD4-BUV563 (GK1.5, BD 612923, 1:500); CD38-BUV395 (clone 90, BD 740245, 1:500); CD38-BUV563 (90/CD38, BD 741271, 1:400); CD80-BV421 (16-10A1, BioLegend 104725, 1:300); CD86-BV605 (GL-1, BioLegend 105037, 1:200); CD138-BUV737 (281-2, BD 624230, 1:500); CXCR4-biotin (BD 551968, 1:200); CXCR4-PE (2B11, Invitrogen 12-9991-82, 1:100); Ephrin-B1-biotin (R&D Systems, BAF473, 1:100); FAS-BUV805 (Jo2, BD 741968, 1:100); FAS-PE-Cy7 (Jo2, BD 557653, 1:500); GL7-FITC (BD 553666, 1:500); GL7-PerCP-Cy5.5 (GL7, BioLegend 144610, 1:500); IgD-BV510 (11-26c.2a, BD 563110, 1:500); IgD-PerCP (11-26c.2a, BioLegend 405709, 1:500); IgG1-BV650 (A85-1, BD 740478, 1:500); IgM-BUV661 (II/41, BD 750660, 1:300); IgM-BUV805 (II/41, BD 749307, 1:500); kappa-APC-Cy7 (1:200); lambda-BV421 (1:400); PD-L2-BUV395 (TY25, BD 565102, 1:75). Secondary detection reagents: SA-APC-Cy7 (BioLegend 405208, 1:100) and SA-BV605 (BioLegend 405229, 1:200). NP-PE (N-5070-1) was used at a working concentration of 5 mg/mL.

### Single-cell multiome (RNA + ATAC) sequencing

Paired scRNA-seq and scATAC-seq libraries were generated with the 10x Genomics Chromium Single Cell Multiome ATAC + Gene Expression kit (10x Genomics; 1000230, 1000283, 1000494, 1000215, 1000212), following the manufacturer’s instructions. In brief, GC B cells were FACS-purified from splenocytes using the antibody panels described above (see Flow Cytometry), washed in PBS containing 0.04% BSA, and 100,000 sorted cells were processed for nuclei isolation per the manufacturer’s protocol, using a hybrid of the standard and low-input procedures. Approximately 20,000 resulting nuclei were loaded onto a 10x Chromium X instrument, and downstream library construction was performed as specified by the manufacturer. Final libraries were assessed for quality on an Agilent Bioanalyzer, quantified with a KAPA Library Quantification Kit, and sequenced on an Illumina NovaSeq.

### Inclusion and ethics statement

All murine studies described here were performed under protocols #2011-0031 and #2025-0008 (PI: Barisic), approved by the Institutional Animal Care and Use Committee (IACUC) at Weill Cornell Medicine, and were carried out in accordance with institutional policies and the Guide for the Care and Use of Laboratory Animals. Each experiment included both male and female animals in age- and sex-matched cohorts; no sex-dependent effects were observed.

### Biological materials

Commercially sourced reagents and previously reported materials are cited in the relevant sections of the text. Novel biological resources generated as part of this work, namely Arid1a^flox/flox^/Ctcf^flox/flox^;Cgamma1-Cre double conditional knockout mouse model and Arid1a^flox/flox^/Stat3^flox/flox^;Cgamma1-Cre, can be obtained from the corresponding authors upon reasonable request, potentially subject to a Material Transfer Agreement.

### scATAC data processing

We used ArchR^34^ to analyze our scATAC-seq data. We inputted the fragment files of our seven GC samples: three WTs, one *Arid1aHet*, one *CtcfHet*, and two *Arid1aHom*. ArchR tiles the genome with the resolution of 500bp and then calculates the highly variable tiles. We used 25000 highly variable genes and LSI dimensionality reduction to bring the dimension to 30. We then used these clusters to create pseudo-bulk replicates and run the peak calling algorithm by MACS2^35^.

### Motif matrices from chromVAR

After getting the peak matrix from ArchR, we ran chromVAR^26^ to get the TF motif accessibility z scores in each single cell. We then smoothed these z scores over 15 nearest cells on the KNN graph derived from LSI dimensions. We used the Vierstra motif set^36^ in chromVAR. This yielded the motif matrix for ~2100 motifs. To filter the motifs with non/low-expressed TFs, we only kept the motifs whose TFs are expressed in at least 500 cells. As the TF motifs could be degenerate and multiple motifs could correspond to one TF, we only kept one motif per TF by choosing the pair that has the maximum Pearson correlation between the motif accessibility scores and RNA expression values. At the end, we ended up with a motif matrix for ~200 motifs which was used as an input for the DynaVelo model.

### scRNA data processing

We used Scanpy^37^ to read the CellRanger outputs for all seven samples. We filtered the cells by keeping the ones with minimum gene count of 200, minimum UMI count of 1000, maximum UMI count of 10000, and percentage of mitochondrial genes less than 25%. We next did total count normalization to the median value of all cells and log1p-normalized (log(x+1)) them. Finally, we kept 2000 highly variable genes to be used in the DynaVelo model.

### Spliced and unspliced counts

We ran velocyto^15^ to read the CellRanger outputs and get the spliced and unspliced counts through loom files for each sample. We used Scanpy to read these loom files and merged the spliced and unspliced matrices to the RNA anndata, ready to be used by scVelo for RNA velocity estimates.

### Estimated RNA velocities by scVelo

We used scVelo to get the estimated RNA velocities. Since cells in each sample might have different dynamics, we ran scVelo per sample. scVelo needs sufficient spliced and unspliced counts to have a reliable RNA velocity estimation. We followed the scVelo pipeline and kept the genes that have more than 10 shared spliced and unspliced counts. This filtering step usually got rid of ~20,000 genes. We normalized the count, spliced, and unspliced matrices based on the median values and log1p-normalized the count matrix. When running scVelo, we specified considering all the genes not only the highly variable ones. The reason for this choice was to have access to maximum number of velocity genes, the genes for which scVelo provides RNA velocity estimates. Depending on the sample, the number of velocity genes was in the range of ~400 to ~1000 genes.

### Genes used in DynaVelo models

We chose four subsets of genes to be used in the DynaVelo models: highly variable genes (2000 genes), velocity genes (~400-1000 genes), expressed TFs (~200 genes), and genes of interest (~20 genes). Highly variable genes carry more information about the cell state transitions. Velocity genes provide additional directional information to guide the learned dynamics. We also included the TFs that were expressed in at least 500 cells to be included in the GRNs as well as the genes of interest in GCs such as *Arid1a* which is an important epigenetic regulator often mutated in the lymphomas. We used the union of these four subsets (~3000-4000 genes) as our modeled genes.

### DynaVelo model details

chromVAR assigns cell-wise TF motif accessibility scores by genome-wide scanning of accessible peaks for TF motifs. Given the crucial role of TFs in regulating target genes within dynamic systems, we hypothesize that integrating TF motif accessibility scores could enhance the understanding of cellular dynamics across transcriptomic and epigenomic spaces. If we denote log-normalized RNA expression values of *m* genes in a cell at the latent time *t* by *x*(*t*) ∈ *R*^*m*^, and chromVAR z-scores of *k* TFs in the same cell by *y*(*t*) ∈ *R*^*k*^, we model the cellular dynamics by a joint ODE as:

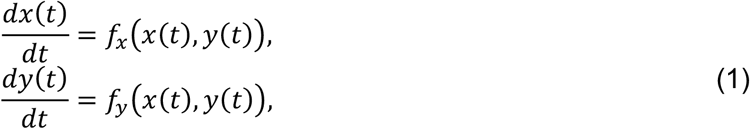

where *f*_*x*_ and *f*_*y*_ are functions that model the dynamics (velocities) of RNA expression and TF motif accessibility, respectively. Here, we have full information of *x*(*t*) and *y*(*t*) and partial information of RNA velocities *dx*(*t*)/*dt* (for “velocity genes”, for which a method like scVelo provides an estimate), and we want to learn the dynamics functions *f*_*x*_ and *f*_*y*_ and the latent times, *t*, of the cells. After learning the dynamics functions, we will recover the RNA velocities of all genes (including non-velocity genes) and define the TF motif velocities *dy*(*t*)/*dt* to reveal important information regarding the dynamics of the chromatin landscape. Instead of working with ODEs in the original high-dimensional spaces, we define an equivalent joint ODE in low-dimensional latent spaces 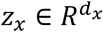 and 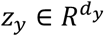, and then map them to the original spaces. We set the default values *d*_*x*_ = *d*_*y*_ = 50 in the models. The rationale behind this is two-fold. First, we would like to have velocity-aware latent spaces that will be informative in illustrating dynamical patterns of the data. Second, since the high-dimensional data are noisy and sparse, learning the dynamics functions in the original spaces is computationally costly and less robust. We define the joint ODE in the latent space as:

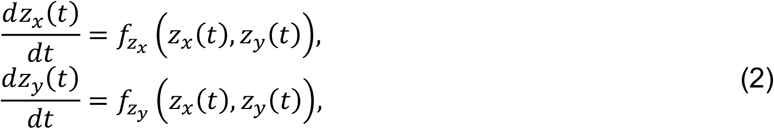

where *z*_*x*_(*t*) and *z*_*y*_(*t*) are respectively the latent space of RNA expression and TF motif accessibility of a cell at the latent time *t*, and 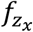 and 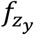 are respectively the dynamics functions of *z*_*x*_ and *z*_*y*_. If we denote the concatenations of these vectors by *z*(*t*) = [*z*_*x*_(*t*), *z*_*y*_(*t*)] and 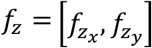, we can write the joint latent ODE as 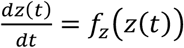, which has the solution 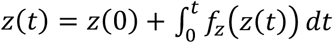 and is known as the initial value problem (IVP). Since we do not know the functional form of *f*_*z*_, we define it as a neural network, which leads to a neural ODE introduced in the seminal work^38^. Neural ODEs are very powerful and efficient models for learning dynamics. After learning the joint cellular dynamics in the latent space, we map them back to the original spaces of RNA expression and motif accessibility by conditional distributions:

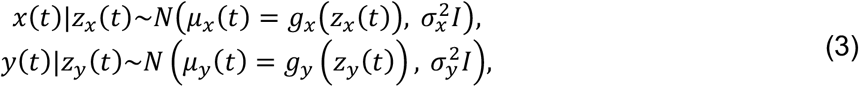

where *g*_*x*_ and *g*_*y*_ are two neural network decoders mapping the latent space to the original spaces, and 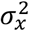 and 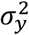 are the known variances of the Gaussian distributions (we assume 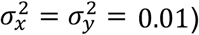. We define the RNA and motif velocities as the time derivatives of *μ*_*x*_(*t*) and *μ*_*y*_(*t*):

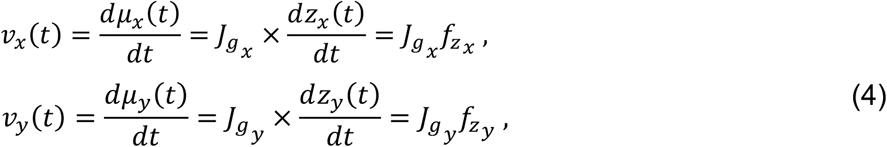

where 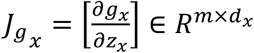 and 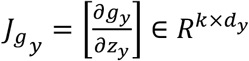 are respectively the Jacobian matrices of the decoder functions *g*_*x*_ and *g*_*y*_. We assume that the initial points of cells are unknown in order to have a general framework for less studied and more complex biological systems such as GC B cells. Therefore, we assume that *z*(0) is a random variable with the prior standard Gaussian distribution *z*(0)~*N*(0, *I*). We also assume that the latent times of cells *t* follow a prior Beta distribution *t*~*Beta*(2,2), as the times should be positive. This leads to the expectation *E*[*t*] = 1 and variance *Var*[*t*] = 0.05. As such, the problem is to learn the posterior probability distributions of *z*(0) and *t* for each cell, the dynamics function *f*_*z*_, and the decoder functions *g*_*x*_ and *g*_*y*_. To this end, we use variational inference techniques by first assuming approximate posterior distributions for *z*(0) and *t* and then using inference networks (or encoders) to learn the parameters of those approximate posteriors, as shown in **Fig. 1a**. We assume the same family of distributions for the approximate posteriors as the priors, that is, 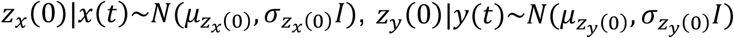, and *t*|*x*(*t*), *y*(*t*)~*Beta*(*α*_*t*_, *β*_*t*_), where 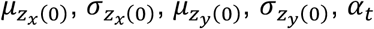, and *β*_*t*_ are encoder neural networks that take in the inputs *x*(*t*) and *y*(*t*) and output the corresponding parameters of the approximate posterior distributions.

In addition to reconstruction loss, we use a KL divergence loss function to ensure that the learned initial (latent time *t* = 0) latent states and cell-specific latent times do not greatly diverge from a standard normal distribution and a Beta distribution, respectively. There are two additional loss functions used in DynaVelo. The first one is a cosine similarity loss function for the RNA velocities, where any reliable RNA velocity method, like scVelo, is used to provide velocity estimates for a subset of genes and supervise the corresponding learned RNA velocities. The second loss function is a velocity consistency loss to ensure that RNA and TF motif velocities lead to similar trajectories of cells in their respective latent spaces. The total loss function has the following form:

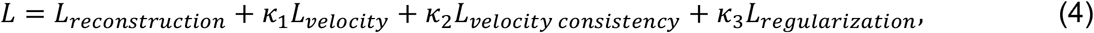

where the first term is the reconstruction loss of RNA expression and TF motif accessibility which maximizes the log-likelihoods:

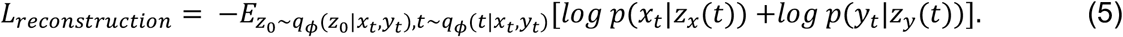

The second term is RNA velocity loss which maximizes the cosine similarity of predicted velocities and scVelo velocities for the velocity genes:

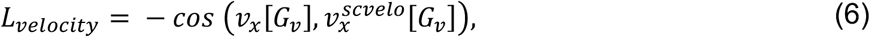

where *G*_*v*_ and 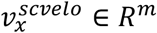 denote respectively the indices of velocity genes as reported by scVelo and the estimated RNA velocities by scVelo. The third term is velocity consistency loss which ensures that cells in two RNA and motif spaces follow consistent trajectories:

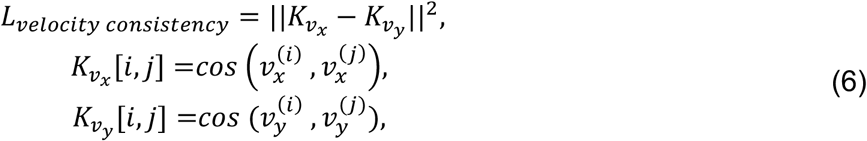

where 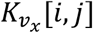 and 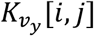 store respectively the cosine similarities of RNA and motif velocity vectors of the cells *i* and *j*. The velocity consistency loss minimizes the distance between cosine similarity of cells in both RNA and motif spaces and thereby helps retain consistent trajectories in both spaces. The fourth term is regularization loss which minimizes the Kullback–Leibler (KL) divergence between the approximate posterior and prior distributions of *z*(0) and *t*:

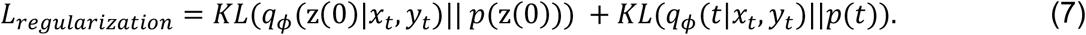

### Hyperparameter selection

Three important hyperparameters of DynaVelo are *κ*_1_, *κ*_2_, and *κ*_3_, which control the strength of different components of the loss function. Depending on how much we trust scVelo velocities, we adjust *κ*_1_, the coefficient of the loss term that supervises the predicted RNA velocities to be consistent with the scVelo velocities for the velocity genes. We advise first making sure that scVelo velocities overall make sense in the biological system to be modeled, which requires domain expertise. We would not recommend supervising DynaVelo velocities too strongly on scVelo if they do not capture the expected trajectories. Since we have expertise in germinal centers (GCs), we made sure that scVelo velocities were capturing the main trajectories of GC B cells. We used different values of *κ*_1_ (1000 and 10000) during training and found that 10000 was a good default value for this hyperparameter in the GC system.

The second hyperparameter, *κ*_2_, controls how consistent the cell trajectories should be in RNA expression and motif accessibility space. If we choose a small value for *κ*_2_, the trajectories might deviate too much between the two spaces, which is not a desired behavior. Therefore, we set *κ*_2_ to the same value as *κ*_1_ (10000 as the default value) to make the trajectories consistent.

Finally, *κ*_3_ controls how close the learned initial point z(0) and latent time *t* of cells should be to their prior distributions, namely *N*(0, *I*) for z(0)and *Beta*(2,2) for *t*. We found that *κ*_3_ is very important in smoothing the learned latent times. If *κ*_3_ is high, it forces all of the cells to start from a similar initial point z(0), and *t* becomes the only source of variation across cells, leading to smooth latent times aligned with the trajectories. Although this is satisfactory for learning the latent times, it might sacrifice the accuracy of predicted delta velocities upon *in silico* perturbations of genes. However, if *κ*_3_ is low, the cells have more freedom to start from any initial point z(0), which might help improve the accuracy of perturbation prediction, while leading to less smooth latent times that do not align well with the trajectories. Therefore, we believe that there is not a universal *κ*_3_ that leads to the best performance in all downstream tasks; rather, this hyperparameter should be selected based on the task. We tested different values for *κ*_3_ (1, 10, 100, 1000) and found 1000 to be a good default for learning velocities and latent times, and 1 to be a good value for perturbation prediction tasks.

### Generating trajectories using DynaVelo

Since DynaVelo is a generative model, it can be used to generate cell trajectories. Given any cell {*x*(*t*), *y*(*t*)}, DynaVelo first maps it to the initial latent state *z*(0) = [*z*_*x*_(0), *z*_*y*_(0)] and finds its latent time *t*, which is simply the mean of the posterior Beta distribution: 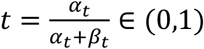. For generating trajectories, we start from *z*(0) at time *t* = 0 and solve the ODEs up to the time *t* = 1 with the step size of *t*_s*tep*_ = 0.01:

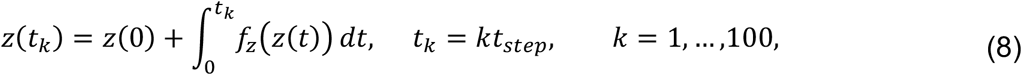

which generates a latent trajectory *T*_*z*_ = [*z*(0), *z*(*t*_1_),, …, *z*(*t*_100_)] with the length of 101 (including the initial point). We then map this latent trajectory to the observational spaces of RNA expression and motif accessibility using their corresponding decoders *μ*_*x*_(*t*) = *g*_*x*_(*z*_*x*_(*t*)) and *μ*_*y*_(*t*) = *g*_*y*_ (*z*_*y*_(*t*)) to generate the trajectories *T*_*x*_ = [*μ*_*x*_(0), *μ*_*x*_(*t*_1_),, …, *μ*_*x*_(*t*_100_)] and *T*_*y*_ = [*μ*_*y*_(0), *μ*_*y*_(*t*_1_), …, *μ*_*y*_(*t*_100_)]. To quantify the uncertainty of these trajectories, we sample *N* = 50 initial latent states from the learned posterior distributions 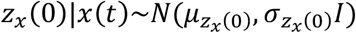 and 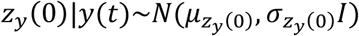 and generate *N* latent trajectories 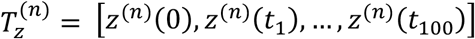 and *N* corresponding RNA expression and motif accessibility trajectories 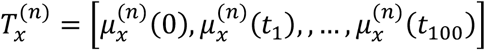 and 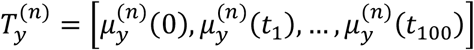 for *n* = 1, …, *N*. We then calculate the mean and standard error of the mean at each time point of the RNA trajectories as (the calculation is the same for motif accessibility trajectories):

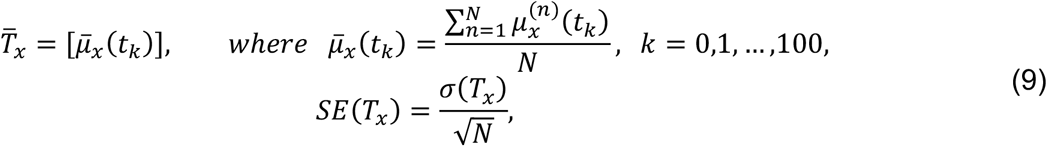

where *σ*(*T*_*x*_) is the standard deviation at teach time point:

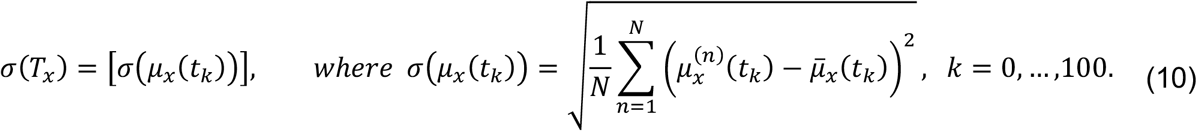

Finally, we calculate the 95% confidence interval of the mean trajectory as:

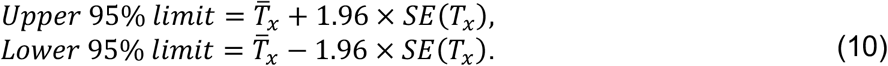

### Calculating the Jacobian matrices for dynamic GRNs

We can calculate the Jacobian matrices, evaluated at individual cells, that represent the derivatives of the RNA and TF motif velocity outputs with respect to the RNA expression and TF motif accessibility inputs. This results in four Jacobian matrices in each cell showing how: (1) RNA expression affects RNA velocity; (2) RNA expression affects motif velocity; (3) TF motif accessibility affects RNA velocity; and (4) TF motif accessibility affects motif velocity. Specifically, once we have learned the velocity functions *f*_*x*_ and *f*_*y*_ in the joint ODE model, we can define four Jacobian matrices for the cell *c* as:

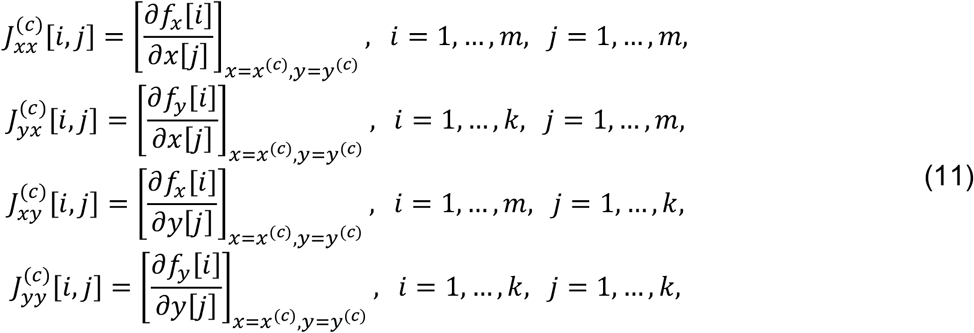

for *c* = 1, …, *n*_*cells*_, where *J*_*xx*_ and *J*_*yx*_ show how the RNA expression of a gene can affect all other RNA and motif velocities, respectively, and *J*_*xy*_ and *J*_*yy*_ show how the motif accessibility of a TF can affect all other RNA and motif velocities. This allows us to understand the gene regulation in a dynamic manner as these Jacobians are cell specific. Since, all the reported GRNs in the literature are static rather than dynamic, we can calculate summary statistics of these Jacobian matrices in any cell states to get the static cell-state-specific GRNs. For example, we can get the transcriptional GRNs in the cell state *S* as:

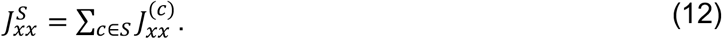

To calculate the Jacobians, we use the definition of partial derivatives:

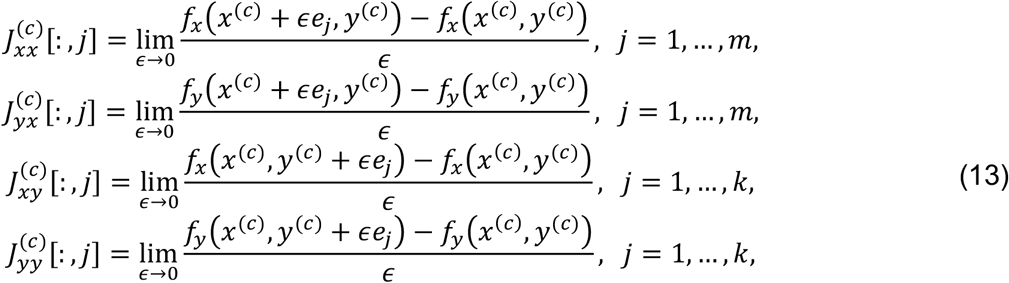

for *c* = 1, …, *n*_*cells*_, where *e*_*j*_ is the unit vector of *j*-th element in either *x*^(*c*)^ or *y*^(*c*)^. We use a small value *ϵ* = 10^−4^ to estimate these Jacobian matrices.

### In silico perturbations

To do *in silico* perturbation of *i*-th non-TF gene, we set its RNA expression to the minimum value over the population of all cells and calculate the change in both RNA and motif velocity vectors (delta velocity) as:

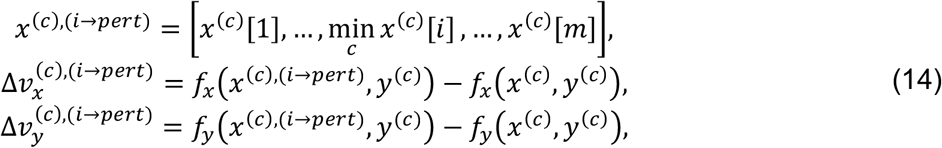

for *c* = 1, …, *n*_*cells*_. To do *in silico* perturbation of *i*-th TF, we set both its RNA expression and motif accessibility to the minimum value over the population of all cells and calculate the delta velocities as (if the *i*-th element in both *x* and *y* vectors correspond to the same TF):

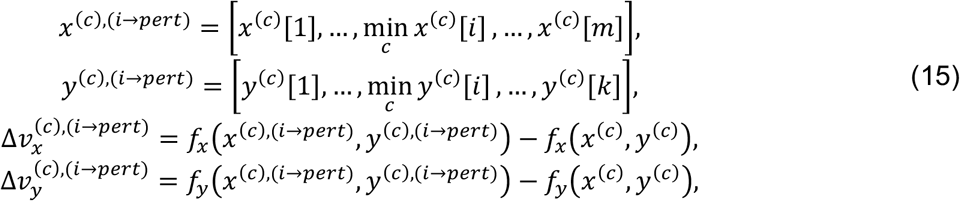

for *c* = 1, …, *n*_*cells*_. These delta velocities could be projected to any lower dimensional spaces such as UMAP to see what the behavior of cells would be and how their trajectories would change after any perturbation. If we look carefully, we can see that these delta velocities are similar to the Jacobian matrices in that they measure the change in velocity functions after any perturbations. The main difference is that the perturbations are infinitesimal in Jacobians but finite in delta velocities. As a result, we can build two delta velocity matrices:

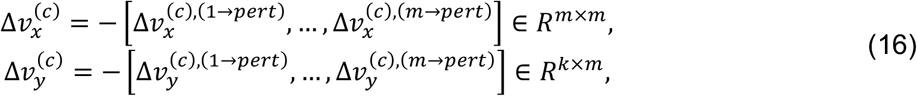

to be used for the dynamic GRN inference. The reason for the negative sign is because the perturbations were knockout (setting to minimum values) instead of over-expression (setting to maximum values), where there is no need for the negative sign. These delta velocities can be used to understand both transcriptional 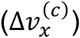 and epigenetic regulation 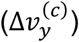 in each cell *c* = 1, …, *n*_*cells*_, or any cell sate *S*_*j*_:

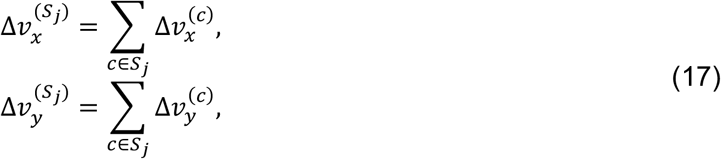

for *j* = 1, …, *n*_*states*_, where *n*_*states*_ is the number of cell states/types. We should note that the dynamic GRNs derived by Jacobians and delta velocities might not be identical due to the nature of perturbations performed in each approach.

### Calculating predicted rescue rates of RNA and motif velocities

Here we describe the details of how we calculate the predicted rescue rates of RNA and motif velocities after any TF perturbation in **Fig. 6d**. Let 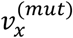 and 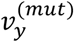 be the RNA and motif velocity vectors of the mutant sample (*Arid1aHet*) and 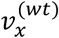 and 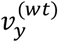 the RNA and motif velocity vectors of the WT sample. We do *in silico* perturbation of the TF *i* in the mutant DynaVelo model and perform a one-sample t-test on the delta velocity vectors in each cell type 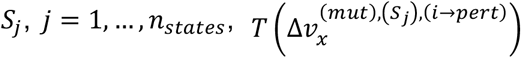 and 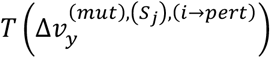, to test if the mean of delta velocities are significantly positive or negative, meaning if this perturbation increases or decreases the velocities in cell type *S*_*j*_. To see if the direction of change in velocities after *in silico* perturbation matches the direction towards the WT sample, we perform another two-sample t-test between the velocity vectors of WT and mutant samples in cell type *S*, i.e. 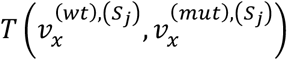 and 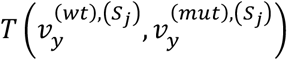. Then we get the significant differntial velocity (DV) genes and motifs from 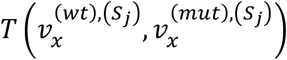 and 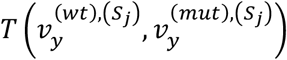 by keeping the ones with *p* < 0.05. Now we calculate the rescue rates of RNA and motif velocities as the fraction of significant DV genes and motifs that match the sign between 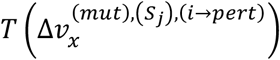 and 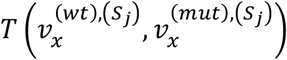 and between 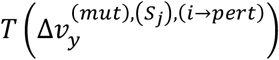 and 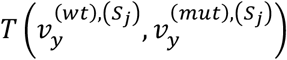 in each cell types *S*. Then, we calcualte the mean rescue rates over all the celltypes, as plotted in **Fig. 6d**. In a nutshell, this shows how much a TF perturbation in a mutant sample can shift its RNA and motif velocities towards the RNA and motif velocities of the WT sample.

### Calculating diffential velocity overlap and direction agreement

We can perform an *in silico* perturbation of the gene *i* in the WT sample, predict the delta velocities, and check how much they shift towards the velocities of the mutant sample, as shown in **Fig. 6b** for the perturbation of *Arid1a*. To this end, we perform the one-sample t-tests 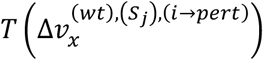 and 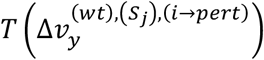 and two-sample t-tests 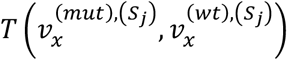 and 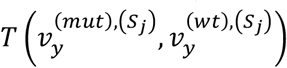 in cell type *S, j* = 1, …, *n*_*states*_. Now we define the DV overlap as the number of overlapped significant DV genes and motifs (*p*_*adj*_ < 0.05) between the one-sample and two-samle tests over the number of genes and motifs. Then we define the DV direction agreement as the number of overlapped significant DV genes for which 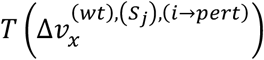 and 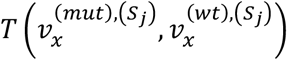 match the sign in each *S*, and overlapped significant DV motifs for which 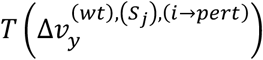 and 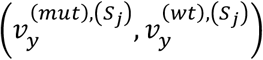 match the sign in each *S*, over th number of DV overlap genes and motifs, respectively. Higher values of DV overlap and direction agreement mean more correct velocity predictions after *in silico* perturbations. True rescure rates reported in **Fig. 6f** for *Arid1aHet*;*CtcfHet* DKO GC are also based on direction agreements in all four data modalities, RNA expression, RNA velocity, motif accessibility, and motif velocity.

### Model architecture details

DynaVelo is a multimodal variational neural ODE framework that jointly models RNA expression and TF motif accessibility dynamics from paired single-cell multiome data. The model takes as input RNA expression vectors *x* ∈ ℝ^*m*^ and TF motif accessibility vector *x* ∈ ℝ^*k*^, where *m* and *k* denote the numbers of genes and TF motifs, respectively. RNA and TF accessibility are first encoded independently into modality-specific latent spaces using shallow neural encoders consisting of a single fully connected layer followed by GELU activation. Specifically, the RNA encoder maps *x* through a linear layer of size [*m, d*_*x*_] followed by GELU, while the TF accessibility encoder maps *y* through a linear layer of size [*k, d*_*y*_] followed by GELU. Default latent dimensions were *d*_*x*_ = *d*_*y*_ = 50. The posterior distributions of latent initial states *z*_*x*_(0) and *z*_*y*_(0) are parameterized as diagonal Gaussian distributions. Separate linear layers estimate the latent means and variances for each modality, with Softplus activation applied to the variance branches to ensure positivity. Latent initial states are sampled using the reparameterization trick and concatenated into a shared latent state 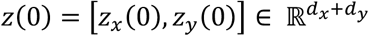.

Latent cellular time is inferred probabilistically using a variational Beta distribution conditioned jointly on RNA and TF accessibility. Separate time encoders, architecturally identical to the expression encoders, map RNA and TF accessibility into latent embeddings of dimensions *d*_*x*_ and *d*_*y*_, respectively. These embeddings are concatenated and passed through two independent linear layers with Softplus activations to infer the Beta distribution parameters *α*_*t*_ and *β*_*t*_. The latent time posterior is therefore defined as *q*(*t*|*x, y*) = *Beta*(*α*_*t*_, *β*_*t*_), with a symmetric Beta prior *Beta*(2,2). During training, latent times are sampled using differentiable reparameterized sampling. Sampled times are sorted before ODE integration to ensure monotonically increasing integration intervals and numerical stability.

Latent dynamics are governed by a shared neural ODE defined on the concatenated latent state. The default ODE function consists of two parallel subnetworks independently predicting RNA latent velocities and TF motif latent velocities. Each subnetwork contains a fully connected layer mapping *d*_*x*_ + *d*_*y*_ → *h*, followed by GELU activation and a second linear layer mapping *h* → *d*_*x*_ or *h* → *d*_*y*_, respectively, where the hidden dimension was typically *h* = 200. The outputs are concatenated to produce the latent velocity field *f*_*z*_(*z*(*t*)). All linear layers in the ODE function were initialized with Gaussian weights (*μ* = 0, *σ* = 0.01) and zero biases. Latent trajectories were solved using the Dormand–Prince adaptive ODE solver (dopri5) from torchdiffeq with adjoint sensitivity optimization enabled during training.

Decoded RNA expression and TF accessibility are reconstructed independently from the latent states. The RNA decoder consists of a single linear layer mapping *d*_*x*_ → *m* followed by ReLU activation to enforce nonnegative reconstructed expression values. The TF accessibility decoder consists of a single linear layer mapping *d*_*y*_ → *k* without output activation, allowing positive and negative chromVAR z-scores. Observed-space velocities were computed by projecting latent velocities through decoder Jacobians using the chain rule. Specifically, decoder Jacobians were computed with automatic differentiation using vectorized Jacobian evaluation (vmap(jacrev)), and observed velocities were obtained as 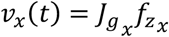 and 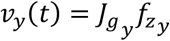, where 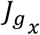 and 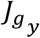 denote the Jacobians of the RNA and TF motif decoders with respect to their latent representations.

The training objective combined reconstruction likelihoods, variational regularization, RNA velocity supervision, and cross-modal velocity consistency. RNA and TF motif accessibility reconstructions were modeled with Gaussian likelihoods having fixed standard deviations 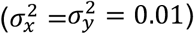. KL divergence penalties regularized both the latent initial states toward a standard normal prior and latent times toward the Beta prior. RNA velocity supervision was implemented using negative cosine similarity between predicted RNA velocities and estimated scVelo RNA velocities, masked to genes with valid velocity estimates. To encourage coordinated multimodal dynamics, a velocity consistency loss minimized the squared difference between pairwise cosine similarity matrices computed from predicted RNA and motif velocities across cells. The total loss was defined as the sum of reconstruction losses, KL penalties, velocity supervision, and consistency regularization, with default weighting coefficients *k*_*t*_ = *k*_*z*0_ = 1000 and *k*_*velocity*_ = *k*_*consistency*_ = 10000. During optimization, cells were additionally weighted inversely proportional to cell-type abundance to mitigate imbalance across populations.

### Model training and inference details

Model optimization was performed using the Adam optimizer with an initial learning rate of 1e−3. For each genotype or sample, cells were randomly partitioned into training and validation sets using a 90%/10% split. The validation set was used for monitoring generalization performance during training. Learning rate scheduling was performed adaptively using a plateau-based scheduler with a multiplicative decay factor of 0.5 and patience of 5 epochs. Early stopping was employed using the validation loss with a patience of 10 epochs, and the model parameters corresponding to the lowest validation loss were retained. Training was performed on a single NVIDIA RTX A6000 GPU (49 GB VRAM). We used batch sizes of 128 and 256 depending on dataset size and memory constraints. Training on datasets containing 5,000–20,000 cells typically required at most 3 hours when using a batch size of 256. We used the maximum epoch size of 200, but early stopping usually got triggered before that.

For probabilistic inference of latent trajectories and velocities after training, the model supports stochastic posterior sampling of both latent initial states and latent times. During prediction, RNA and motif velocities were estimated by repeatedly sampling from the posterior distributions of latent initial state and latent times. Specifically, the model was evaluated *n*_*samples*_ times (default *n*_*samples*_ = 100), generating independent samples of predicted latent states, latent velocities, RNA and motif velocities. Final predicted quantities were obtained by averaging across posterior samples, while posterior uncertainty estimates were computed using the corresponding sample standard deviations. This Monte Carlo procedure reduces sensitivity to stochastic latent sampling and provides more robust velocity estimates.

For Jacobian estimation, stochastic sampling was disabled to obtain deterministic dynamical functions. In this setting, the posterior means of latent initial states and latent times were used directly instead of sampled values. Consequently, Jacobians were computed deterministically from the learned mean latent trajectories, enabling stable estimation of local regulatory interactions and perturbation responses.

In silico perturbation could also be done in both deterministic and stochastic modes. In deterministic mode, perturbation effects were estimated using posterior mean latent variables and latent times, resulting in faster inference. In stochastic mode, perturbation effects were averaged across repeated posterior samples, analogous to velocity estimation. The stochastic perturbation procedure generally provides more robust estimates, although computational cost scales approximately linearly with the number of posterior samples. Users can therefore trade computational efficiency for robustness depending on the application and dataset size. We use the deterministic mode as a default.

## Data availability

We have used seven multiome samples: Ctcf-29 (WT), Arid1a-C3 (WT), Arid1a-D3 (WT), Arid1a-M3 (*Arid1aHet*), Arid1a-M1 (*Arid1aHom*), Arid1a-C2 (*Arid1aHom*), and Ctcf-30 (*CtcfHet*). The first six samples are available on the NCBI Gene Expression Omnibus via GSE237990, and the last sample (*CtcfHet*) via GSE294051. We have also provided the finalized preprocessed AnnData objects at the private Figshare repository https://figshare.com/s/84862d73cc45f47c9d94, which can be used to train the DyneVelo models.

For the sake of biological validations, we have generated six new multiome samples: WT-14 (WT, N=15327), WT-5 (WT, N=11852), HET-21 (*Arid1aHet*, N=14070), HET-553 (*Arid1aHet*, N=15014), DKO-538 (*Arid1aHet;CtcfHet*, N=14314), and DKO-552 (*Arid1aHet;CtcfHet*, N=12460), where N is the number of cells after filtering. These six samples are available on the NCBI Gene Expression Omnibus via GSE334370 and access token “mnizymoohngvnwx”.

## Code availability

https://github.com/karbalayghareh/DynaVelo includes the code package and a tutorial for training DynaVelo models. It also includes Python notebooks to generate the figures of this paper.

## Acknowledgements

A.K. and C.S.L. are funded by IGVF U01 NIH/NHGRI HG012103. D.B. is supported by R00 CA246080. C.R.C. is supported by NIH/NIAMS T32AR071302.

## Author contributions

A.K. developed DynaVelo, performed all the analyses, and co-wrote the paper with C.S.L.; C.R.C. annotated the cell types of the GC B cells; D.B. generated the GC B cell multiome datasets and carried out validation experiments; C.S.L. and A.M. supervised the research; C.R.C., D.B., and A.M. provided biological interpretation of computational results.

## Supplementary Note

### Benchmarking other dynamical models

For the purpose of biological validations, we generated six new multiome GC datasets, including two wildtype samples (WT14 and WT5), two *Arid1aHet* samples (HET21 and HET553), and two *Arid1aHet*;*CtcfHet* double mutant samples (DKO538 and DKO552) with heterozygous loss of *Arid1a* and *Ctcf*. **Fig. S3** shows that DynaVelo’s predicted latent times in all six samples are the most robust, as compared to scVelo and MultiVelo. DynaVelo’s latent times always identify plasmablasts (cyan cluster inside the red circle in **Fig. S3)** as one of the terminal states of the germinal centers, while MultiVelo has difficulty in this task, and its latent times are not reproducible across the samples of the same genotype (WT, *Arid1aHet*, and *Arid1aHet*;*CtcfHet* DKO). Although scVelo could find a subset of the plasmablasts as the terminal state, it also identifies a subset of these cells as the root cells, which is not consistent with the GC biology. Furthermore, DynaVelo can correctly identify the centroblasts (light blue cluster inside the blue circle in **Fig. S3**) as the root cells, while scVelo often assigns a higher latent time for those cells, contradicting GC biology. Overall, these results show that DynaVelo’s latent time predictions are high fidelity and robust across samples.

### Model robustness and reproducibility

To evaluate the robustness and reproducibility, we ran the DynaVelo model using 10 different seeds in the WT GC sample (CtcfWT29). Each seed randomizes both the weight initializations of neural networks and training cell splits. To reduce the computational cost, we sampled and fixed 128 cells from the WT GC cells and computed the following quantities in each of 10 models: RNA velocity (vx), motif velocity (vy), four Jacobian matrices (Jxx, Jxy, Jyx, and Jyy), delta RNA velocity (delta_vx), delta motif velocity (delta_vy), and latent times. Note that we used 26 important GC genes for the perturbations in this analysis. Considering the 2656 genes and 169 TFs used in the models, the tensors have the following shapes. vx: [128, 2625], vy: [128, 169], Jxx: [128, 26, 26], Jxy: [128, 26, 169], Jyx: [128, 169, 26], Jyy: [128, 169, 169], delta_vx: [128, 2625, 26], delta_vy: [128, 169, 26], latent_time: [128]. We then flattened all the tensors and calculated the Pearson correlation between 10 seeds, leading to 45 pairwise correlations.

**Fig. S4** shows the distributions of these 45 correlations for each quantity. We can see that the predicted RNA and motif velocities (vx and vy) and latent times of the cells are highly reproducible across seeds with median correlations of 0.97, 0.94, and 0.99. The predicted Jacobians and delta velocities are also well correlated across the seeds, albeit less so compared to the velocities (0.68, 0.69, 0.6, and 0.55 for Jxx, Jxy, Jyx, and Jyy, respectively; 0.52 and 0.6 for delta_vx and delta_vy, respectively). This highlights an important observation that predicting the effects of perturbations is more difficult. We also noticed that using a bigger *ϵ* for calculating the Jacobians leads to more robust results. We used *ϵ* = 0.1 for this analysis. As explained in the manuscript, *ϵ* is used to approximate the Jacobians by perturbing the input gene expression or TF motif accessibility values by adding *ϵ* to the inputs and measuring the change in RNA and motif velocities in the output of the model.

### Model generalizability

DynaVelo predictions can generalize from dataset A to dataset B (without being trained on dataset B) as long as we expect similar trajectories and cell dynamics. For example, if we have two wildtype (WT) germinal center (GC) samples, we expect them to have roughly the same trajectories, even though there might be subtle variations. However, this assumption does not hold between two different conditions, for example between wildtype and mutant samples. If we do some intervention in the same GC system (such as knocking out a gene like *Arid1a* or *Ctcf*), it might drastically change the underlying dynamics of the cells. That is why we strongly suggest running any dynamic model such as DynaVelo or scVelo on cells from the same condition. Therefore, it is not possible to train the DynaVelo model on the WT samples and use it directly in the mutant GC samples (without performing *in silico* perturbation).

What DynaVelo can do is generalize between datasets under the same biological conditions, so the assumptions of having similar trajectories hold. We tested this in our new GC datasets, generated for the sake of biological validations for the revised manuscript. In the new batch of GC samples, we have two WT GC samples, WT5 and WT14. We first trained DynaVelo only on the sample WT5. Next we trained DynaVelo only on WT14 and applied it to predict the latent times and RNA and motif velocities of the WT5 cells to see how the model generalizes to unseen datasets. **Fig. S5a** shows that the RNA and motif velocities in both cases (WT5→WT5 and WT14→WT5) are very similar. We believe that subtle variations between the samples also make sense, as we cannot expect the two systems to have 100% the same dynamics. **Fig. S5b** shows that the predicted latent times of the cells also are similar in the two cases, where centrocytes, prememory, and plasmablast cells have the highest latent times, matching the biology of WT GCs. To further quantify the consistencies between the two cases, we calculated Pearson correlations and plotted the scatterplots in **Fig. S5c**, where we see correlations of 0.83, 0.75, and 0.76 between the RNA and motif velocities and latent times between the two cases. These results show that DynaVelo can be applied to model dynamics in other datasets without retraining on as long as the biological conditions are the same.

### RNA-only DynaVelo

We should clarify that we have implemented DynaVelo in two different modes. The default mode is for multiome data with RNA and ATAC readouts in the same cells. The second mode is for the scenario that only RNA data is available. Since we have multiome data, we have always used the multiome DynaVelo, where we use TF motif accessibility as additional information that can enrich our modeling of cellular dynamics. Since DynaVelo needs some directional velocity information to guide the dynamics of cells, currently it cannot be robustly used in ATAC-only mode, as there is no equivalent of RNA velocity in the ATAC space. Thanks to the reviewer’s comment, we also tested the RNA-only DynaVelo in our WT GC samples to see how well it performs. **Fig. S6a** shows that RNA-only DynaVelo can also capture the main GC trajectories, which are similar to the trajectories learned by the multiome DynaVelo in the majority of cell types. Regarding the learned latent times, even though the RNA-only model identifies prememory and plasmablast cells as terminal states, it also highlights centrocytes as another potential terminal state, which is incorrect. However, the multiome model only highlights the two correct GC terminal states, prememory and plasmablast cells (**Fig. S6a**). Finally to quantify which model can better predict the RNA velocities in heldout test cells, we calculated the cosine similarity between predicted versus scVelo velocities (serving as ground truth velocities). **Fig. S6b** shows the scatterplots of these cosine similarities in each test cell (~2700), where we can see that multiome and RNA-only models have mean Pearson correlation of 0.7 and 0.67 (Wilcoxon p-value 2.18e-155). These analyses overall show that multiome DynaVelo is better than RNA-only DynaVelo in capturing cellular dynamics and latent times of cells, but RNA-only models can also be used in the single-modality setting, i.e. for scRNA-seq datasets.

**Supplementary Figure S1.**
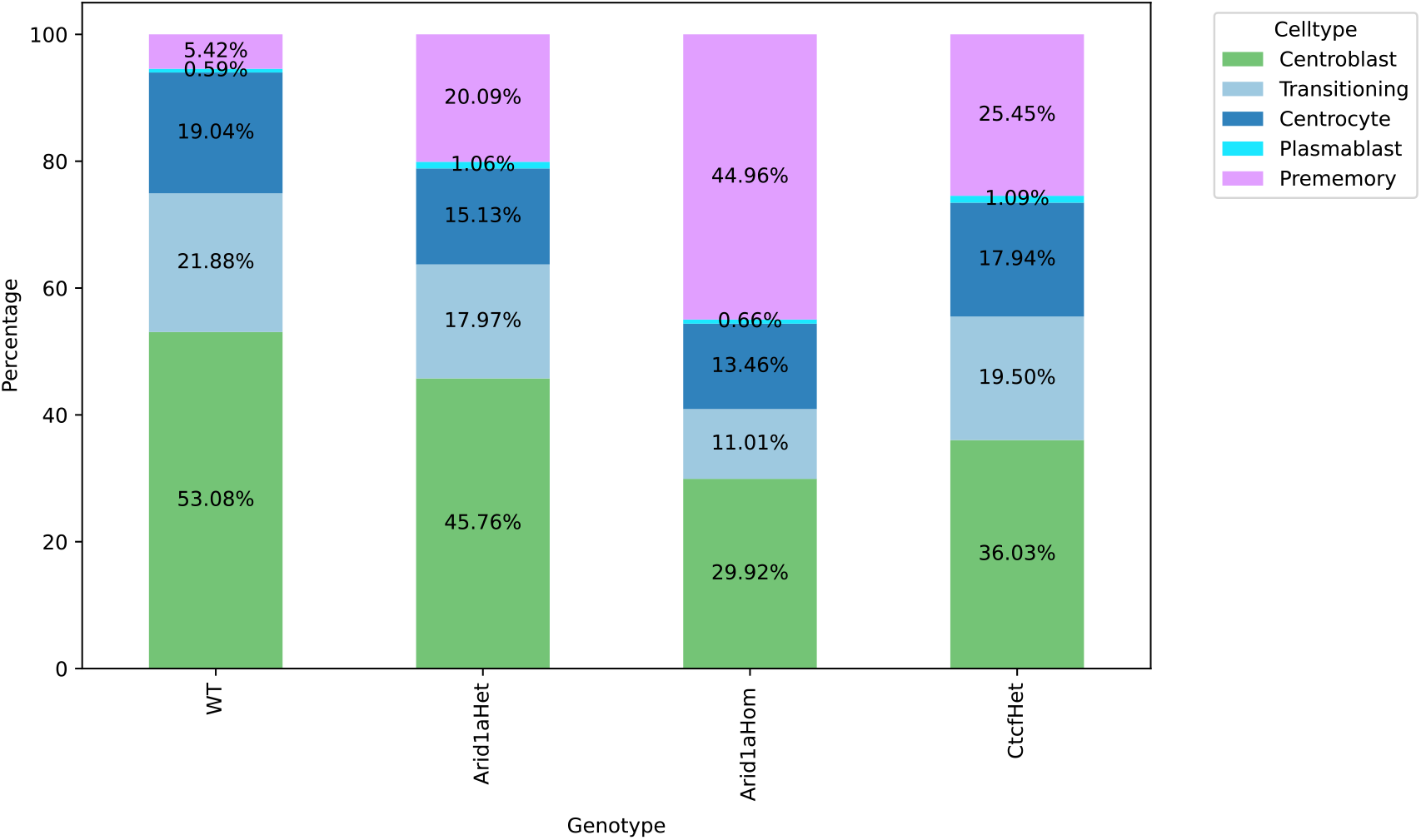
Percentages of cell types in each genotype in mice GC B cells. The percentage of Prememory cells increases in both Arid1a and Ctcf mutant samples.

**Supplementary Figure S2.**
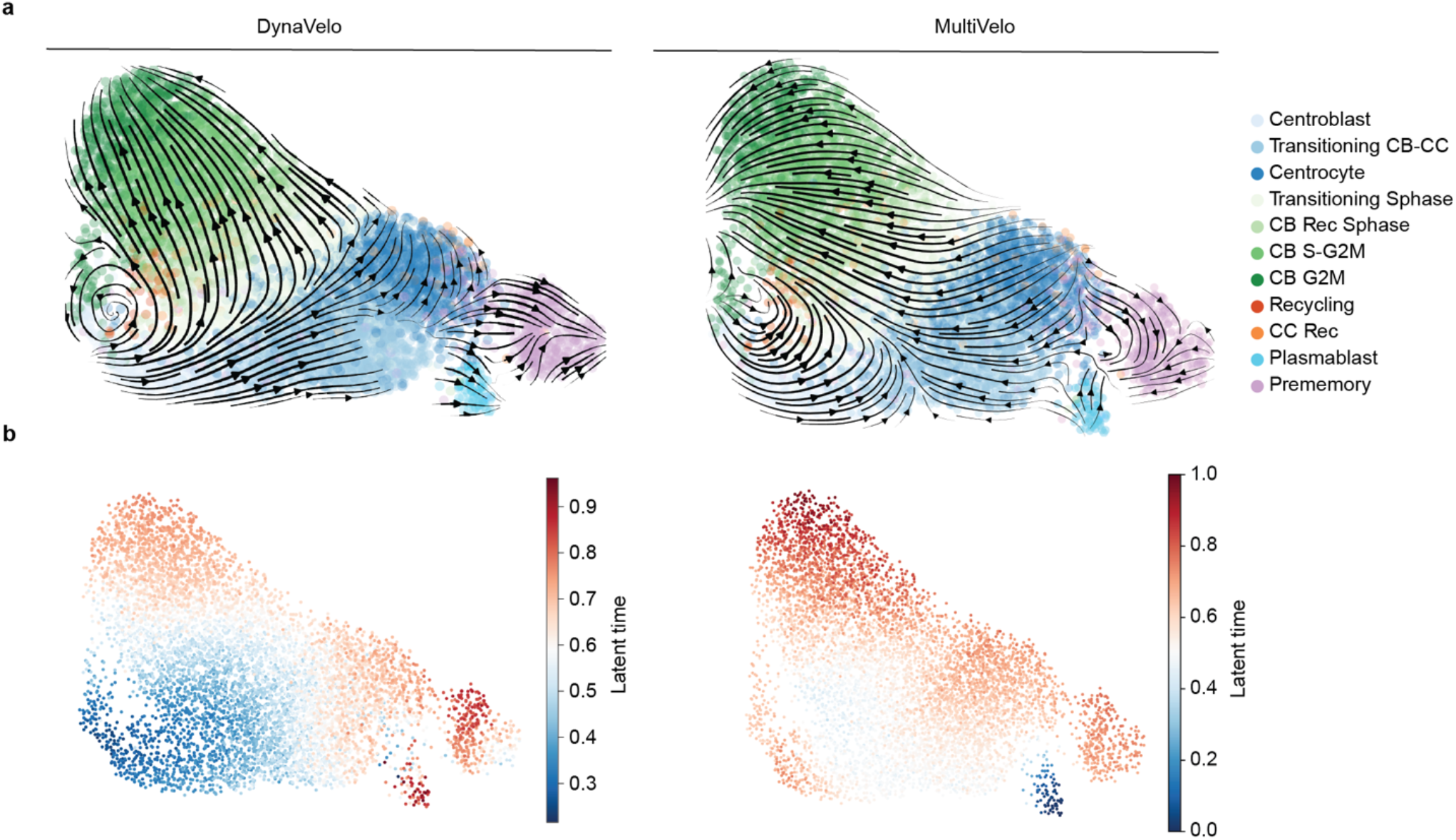
DynaVelo’s RNA velocities and latent times better capture the biology of WT GCB cells than MultiVelo. **a**. The RNA velocities of DynaVelo and MultiVelo projected on the same RNA UMAP space. **b**. The latent times learned from DynaVelo and Multivelo projected on the same RNA UMAP space.

**Supplementary Figure S3.**
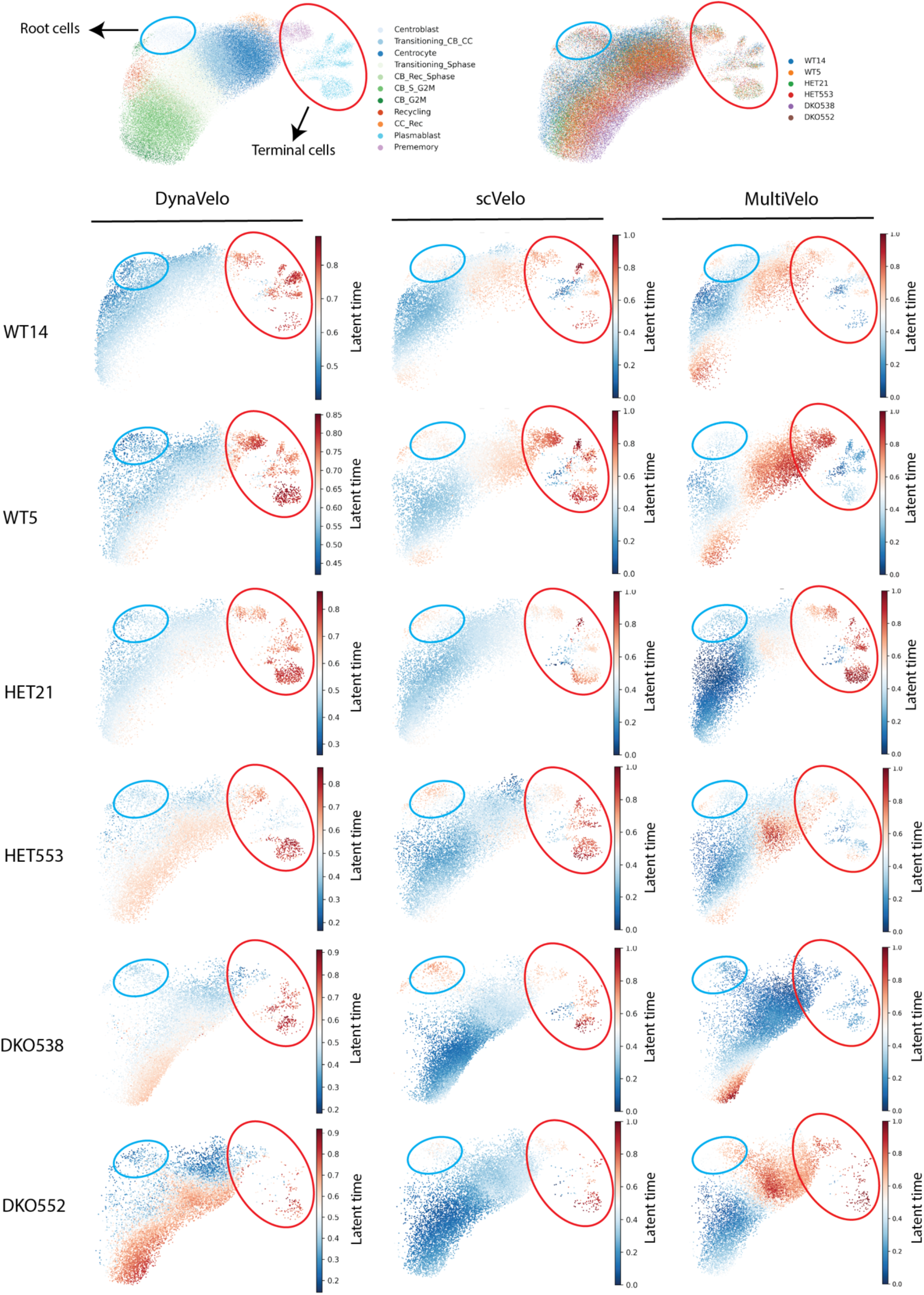
Latent times inferred by different methods in germinal center B cells.

**Supplementary Figure S4.**
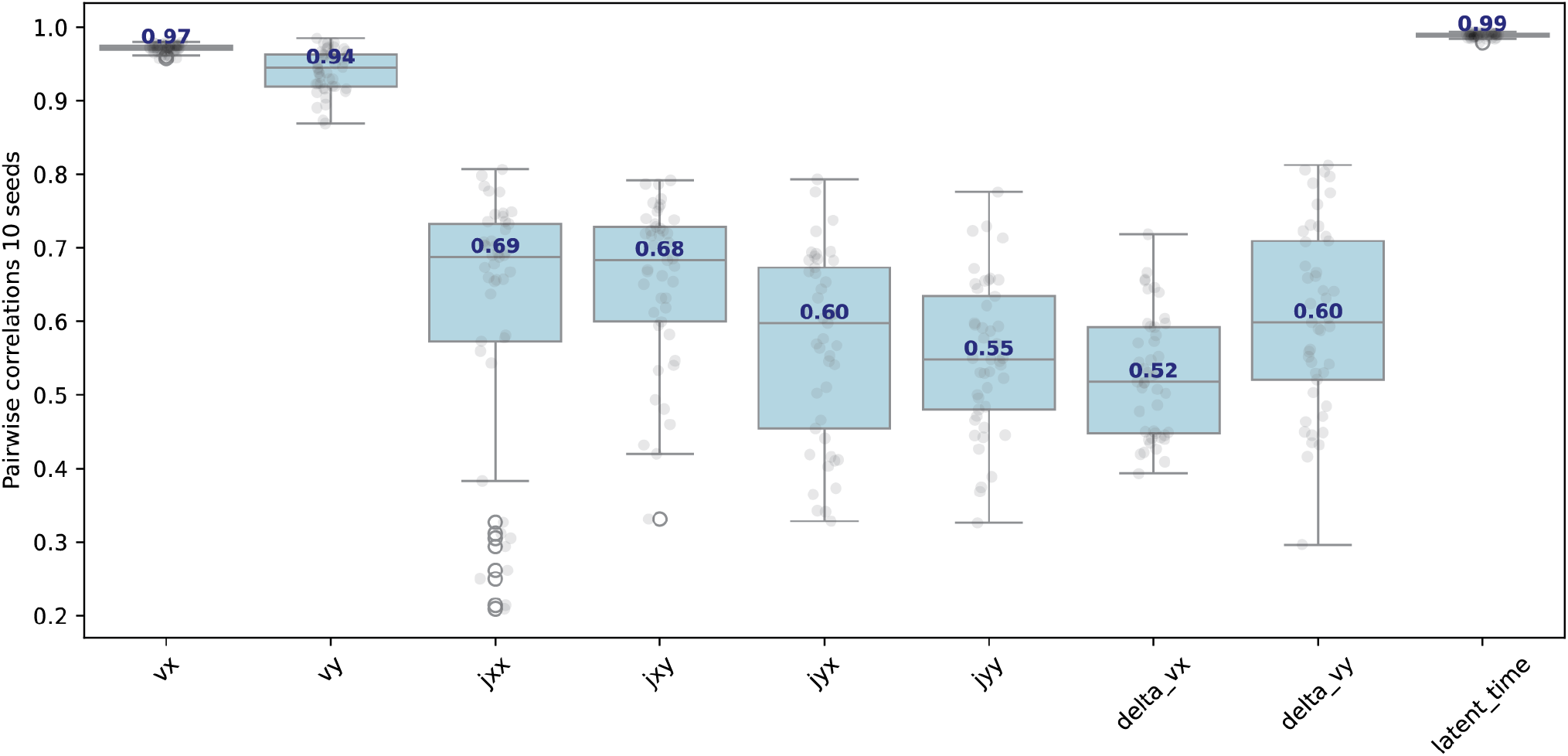
Reproducibility analysis. Pairwise Pearson correlations between 10 random seeds.

**Supplementary Figure S5.**
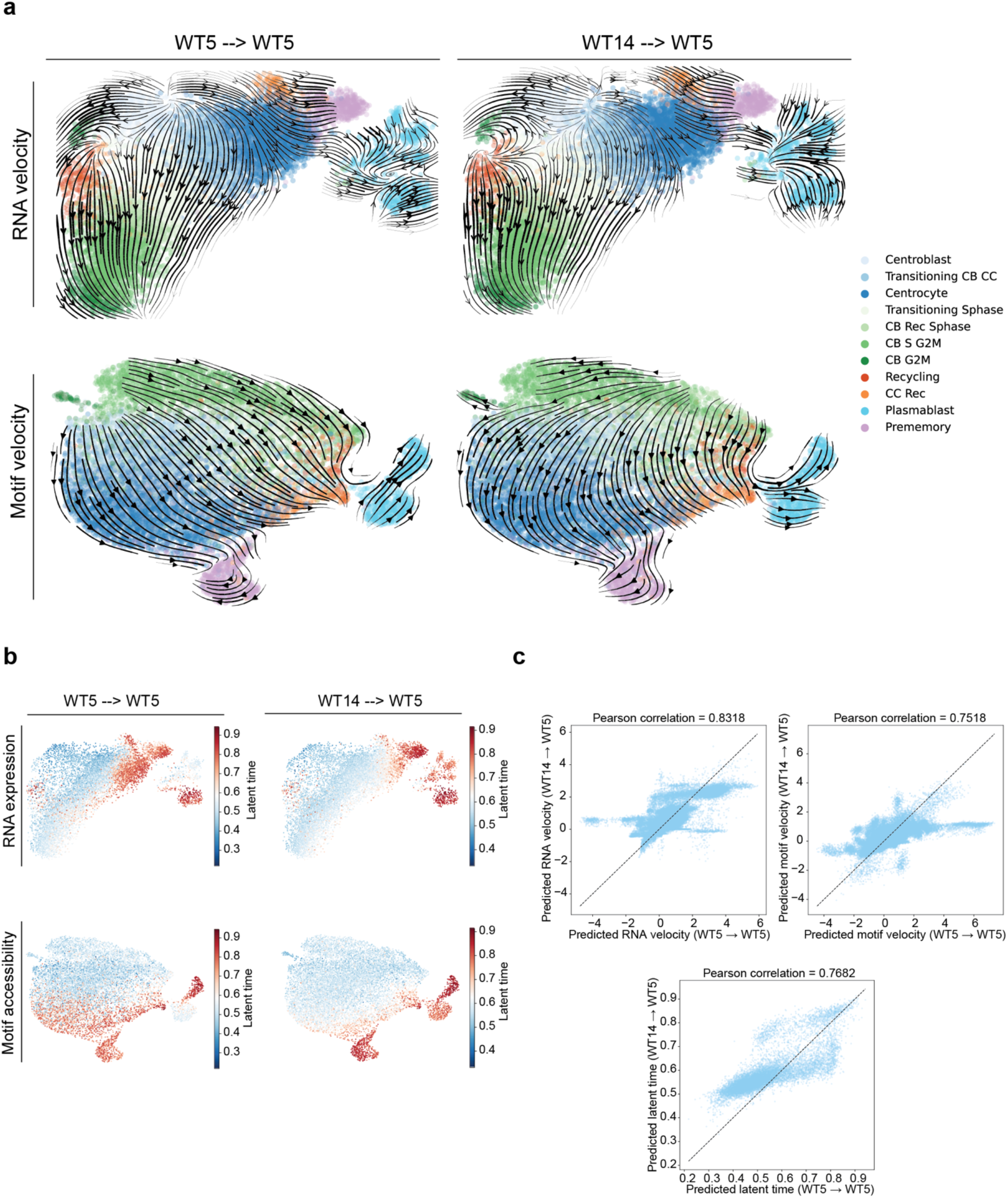
Generalization analysis.

**Supplementary Figure S6.**
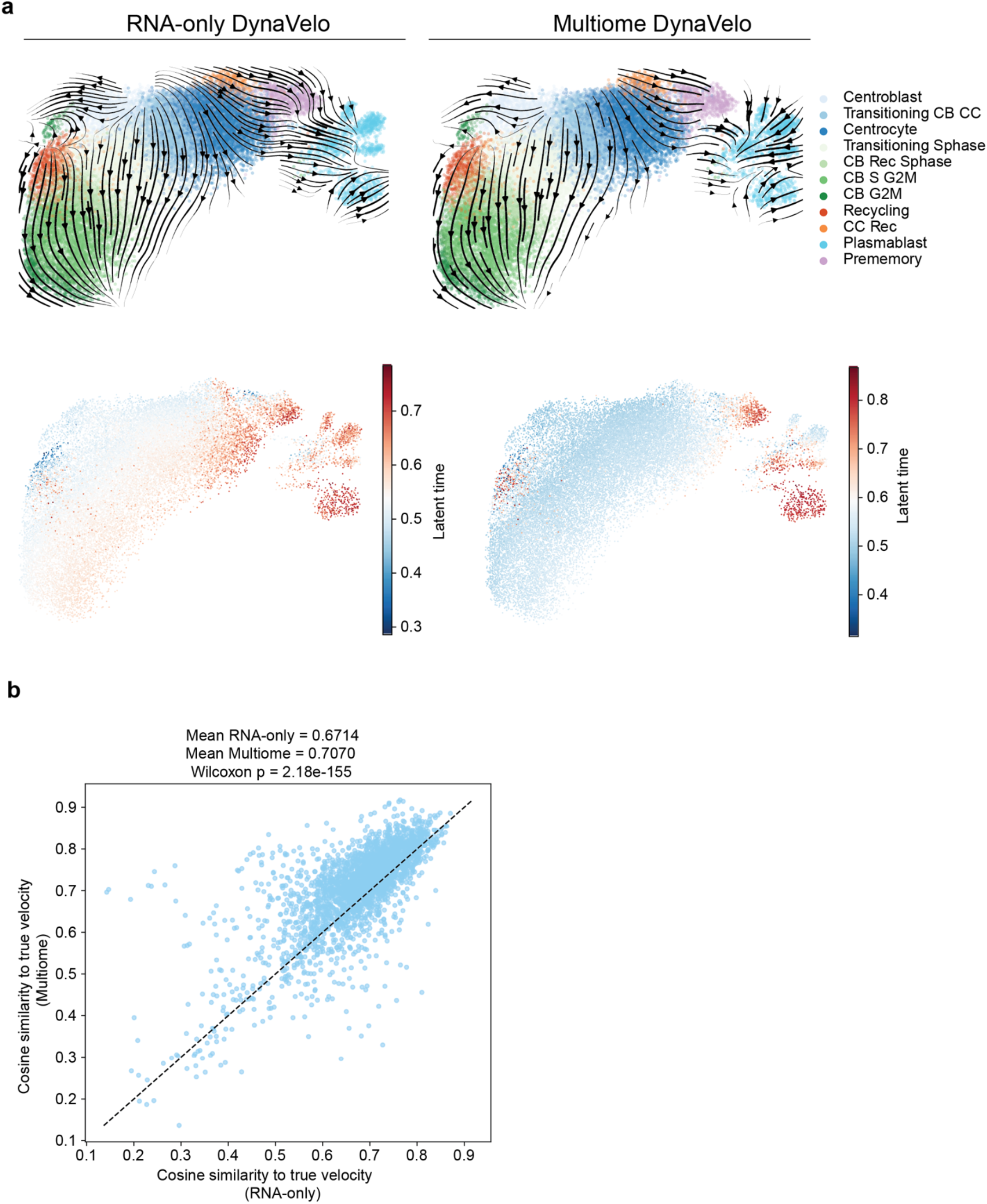
RNA-only vs. multiome DynaVelo.

